# Deciphering the gut microbiome of grass carp through multi-omics approach

**DOI:** 10.1101/2023.03.14.532499

**Authors:** Ming Li, Hui Liang, Hongwei Yang, Qianwen Ding, Rui Xia, Jie Chen, Wenhao Zhou, Yalin Yang, Zhen Zhang, Yuanyuan Yao, Chao Ran, Zhigang Zhou

## Abstract

**Background:** Aquaculture plays an important role in global protein supplies and food security. The ban on antibiotics as feed additive proposes urgent need to develop alternatives. Gut microbiota plays important roles in the metabolism and immunity of fish, and has the potential to give rise to novel green inputs for fish culture. However, our understanding of fish gut microbiome is still lacking.

**Results:** We identified 575,856 non-redundant genes by metagenomic sequencing of the intestinal content samples of grass carp. Taxonomic and functional annotation of the gene catalogue revealed specificity of the gut microbiome of grass carp compared with mammals. Co-occurrence analysis indicated exclusive relations between the genera belonging to Proteobacteria and Fusobacteria/Firmicutes/Bacteroidetes, suggesting two independent ecological groups of the microbiota. The association pattern of Proteobacteria with the gene expression modules of fish gut and liver was consistently opposite to that of Fusobacteria, Firmicutes and Bacteroidetes, implying differential functionality of Proteobacteria and Fusobacteria/Firmicutes/Bacteroidetes. Therefore, the two ecological groups were divided into two functional groups, i.e., Functional Group 1: Proteobacteria; Functional Group 2: Fusobacteria/Firmicutes/Bacteroidetes. Further analysis revealed that the two functional groups differ in genetic capacity for carbohydrate utilization, virulence factors and antibiotic resistance. Finally, we proposed that the ratio of “Functional Group 2/Functional Group 1” can be used as a biomarker that efficiently reflects the structural and functional characteristics of the microbiota of grass carp.

**Conclusions:** The gene catalogue is an important resource for investigating the gut microbiome of grass carp. Multi-omics analysis provides insights into functional implications of the main phyla that comprise the fish microbiota, and shed lights on targets for microbiota regulation.

## 1. Introduction

Fish consumption accounts for 1/6 of the world’s animal protein intake (FAO, 2020). Due to the limited resources of capture fisheries, aquaculture has become the main way to improve the global supply of fish products [1, 2]. The limitations of production factors (land, feed, etc.) and aquaculture environmental stresses (pathogens, parasites, etc.) have resulted in continuous challenges to the high-efficiency and green development of aquaculture [3-5]. Moreover, the ban on antibiotics as feed additive proposes urgent need to develop green inputs as alternatives.

Studies of fish microbiome have the potential to give rise to green inputs for the aquaculture industry [6]. The important role of the commensal microbiota in immune homeostasis, disease and health has been demonstrated in humans [7-9], and a comprehensive gut microbiome gene catalogue was established [10, 11]. Furthermore, metagenomics studies promoted our understanding of the vital functions performed by commensal microbes in livestock and poultry. For example, a metagenomic study in goat constructed 719 high-quality metagenome-assembled genomes (MAGs) and revealed their functions in the production of short-chain fatty acids (SCFAS) (e.g., propionic acid and butyric acid, etc.) [12]. Microbial function was discovered for fibre digestion in the rumen and a potential cross-talk between microbiome and host cells was confirmed in dairy cows by metagenomic sequencing [13]. A gut microbial gene catalogue was constructed in chicken and metagenomic analysis provided insights into the growth-promoting effect of *Macleaya cordata* extract (MCE) [14]. Studies in fish have also found that commensal microbiota plays important roles for the host metabolism and immunity [15]; however, the microbial gene catalogue has not yet been constructed in fish species and functions of main phyla in the aspect of host-interaction remains unclear. Weighted gene co-expression network analysis (WGCNA) has been used in correlations between host gene sets and factors (such as phenotypic traits and environmental factors) [16], which helps to identify key relationships between gene co-expression modules and microbial taxa [17].

Diet is a key factor that influences the structure and function of the gut microbiota [18]. Feeds in aquaculture have been shifting from animal proteins derived from marine resources to plant proteins [4]. Replacement of animal proteins by plant proteins is common in aquaculture, giving rise to formulations with differential percentage of animal versus plant protein sources. Thus, carnivorous (animal protein-dominated), omnivorous (relatively balanced in animal and plant proteins) and herbivorous (plant protein-dominated) diets have become distinctive features of feeds in aquaculture, and microbial alteration associated with the three dietary types is representative when investigating the structural and functional characteristics of fish microbiota.

Grass carp (*Ctenopharyngodon idella*), belonging to the Cyprinidae family, is the most important freshwater farmed fish species in China. At present, the production value of grass carp ranks the first among cultured freshwater fish globally. In this study, we investigate the structural and functional characteristics of the gut microbiome of grass carp. Metagenome sequencing of microbiota was conducted in fish fed carnivorous, omnivorous and herbivorous diets. The gene catalogue was established, and the functional implications of key microbial taxa in the aspect of host-interaction were evaluated by multi-omics approach.

## 2. Materials and methods

### 2.1 Experimental diets

According to the nutritional requirements of NRC (2011) and Wang et al. [15], the experimental formulations of different diets for grass carp were designed with equal nitrogen and lipid levels (Table 1). In brief, soy protein concentrate and wheat gluten protein were the protein sources for the herbivorous diet (HD), and casein and gelatin were the protein sources for the carnivorous diet (CD). Equal proportions of protein sources in the carnivorous and herbivorous diets make up the omnivorous diet (OD). Microcrystalline cellulose has been added as a dietary fiber for herbivorous fish. Soybean oil is applied as a fat source and adapted to the high fat level feeding levels in aquaculture.

**Table.1.** Experimental formulations for different diets of grass carp.

| Ingredient (g/kg diet) | Carnivorous diet | Omnivorous diet | Herbivorous diet |
| --- | --- | --- | --- |
| Soybean protein concentrate | 0 | 210.32 | 420.64 |
| Vital gluten | 0 | 44 | 88 |
| Casein | 304 | 152 | 0 |
| Gelatin | 76 | 38 | 0 |
| Vital gluten | 310 | 265 | 220 |
| Soybean oil | 84.2 | 82.7 | 81.1 |
| Lysine | 0 | 3.69 | 7.37 |
| Methionine | 7.24 | 9.245 | 11.25 |
| VC phosphate | 1 | 1 | 1 |
| Vitamin premix<sup>1</sup> | 2 | 2 | 2 |
| Mineral premix<sup>2</sup> | 2 | 2 | 2 |
| Monocalcium phosphate | 20 | 20 | 20 |
| Choline chloride | 2 | 2 | 2 |
| Microcrystalline cellulose | 80 | 80 | 80 |
| Zeolite powder | 111.56 | 88.045 | 64.64 |
| Total | 1000 | 1000 | 1000 |
| Conventional nutrients |  |  |  |
| Crude protein (%) | 35.27 | 35.25 | 35.95 |
| Crude lipid (%) | 8.47 | 8.27 | 8.64 |
| Crude ash (%) | 4.72 | 4.06 | 3.94 |
| Dry matter (%) | 95.28 | 95.94 | 96.06 |

1. Containing the following (g/kg vitamin premix): thiamine, 0.438; riboflavin, 0.632; pyridoxine, 0.908; d-pantothenic acid, 1.724; nicotinic acid, 4.583; biotin, 0.211; folic acid, 0.549; vitamin B-12, 0.001; inositol, 21.053; menadione sodium bisulfite, 0.889; retinyl acetate, 0.677; cholecalciferol, 0.116.
2. Containing the following (g/kg mineral premix): CoCl<sub>2</sub>·6H<sub>2</sub>O, 0.074; CuSO<sub>4</sub>·5H<sub>2</sub>O, 2.5; FeSO<sub>4</sub>·7H<sub>2</sub>O, 73.2; NaCl, 40.0; MgSO<sub>4</sub>·7H<sub>2</sub>O, 284.0; MnSO<sub>4</sub>·H<sub>2</sub>O, 6.50; KI, 0.68; Na<sub>2</sub>SeO<sub>3</sub>, 0.10; ZnSO<sub>4</sub>·7H<sub>2</sub>O, 131.93; Cellulose, 501.09.

### 2.2 Experimental design and sample collection

Grass carp with an initial weight of about 20 g were selected and maintained in the culture system for 3 weeks. Before the formal experiment, the fish were treated with the CD diet for 1 week. Then, 108 fish of the same-sized fish were randomly allocated to three tanks and fed three times a day by satiety feeding, with one of the three diets CD/OD/HD for three weeks (Supplementary Fig. 1A). During the experiment, the water temperature was maintained at 26±2°C. At week 1 and 3, 15 fish were randomly selected from each dietary group to form six biological replicates, with each replicate consisting of a mixture of two to three fish. The intestinal contents were sampled 4-6 h after feeding, and the corresponding intestinal and liver tissues were also sampled. All samples were quickly frozen directly in liquid nitrogen and finally transferred to -80°C.

### 2.3 DNA extraction and metagenome sequencing

Extraction of DNA from intestinal contents following the protocol described in the kit (Power Soil® DNA Isolation kit). The Illumina NovaSeq6000 sequencing technology platform was used to perform sequencing by shotgun strategy. The detailed experimental procedure was performed according to the standard protocol provided by Illumina, including DNA quality testing, library construction, library quality testing and library sequencing.

### 2.4 RNA extraction, transcriptome sequencing and data analysis

RNA extraction from intestinal and liver tissues was performed by reference to published methods [19]. The extracted RNA was used to construct the library as previously described [19], followed by sequencing using the Illumina NovaSeq6000 technology platform. A total of 466.18 Gb Clean Data was obtained from 72 transcriptome sequencing samples. After the clean data were mapped onto the reference genome [20] by HISAT2 (version 2.0.4,--dta -p 6 --max-intronlen 5000000) [21], the transcript was assembled and using String Tie (Version V1.3.4d,--merge -F 0.1 -T 0.1) [22]. Gene expression was quantified by FPKM method [23]. FoldChange > 1.5 and *p*-value < 0.05 was considered to be differentially expressed genes (DEG) using edgeR (version3.8.6) [24]. Enrichment analysis of KEGG pathways was performed using clusterProfiler (version 3.10.1), and qvalue < 0.05 was considered to be a significant enrichment. Default parameters were used for unlisted parameters.

### 2.5 Metagenomic assembly and non-redundant gene catalogue construction

The Raw Tags were filtered using Trimmomatic (version 0.33) to obtain high quality sequencing data (CleanTags) with the parameters (LEADING:3 TRAILING:3 SLIDINGWINDOW:50:20 MINLEN:100); and the bowtie2 (version 2.2.4) was used to perform sequence alignment with the host genome removing host contamination. Metagenome assembly was performed using the software MEGAHIT (Version 1.1.2), and contig sequences shorter than 300 bp were filtered [25]. The QUAST (Version 2.3) software was used to evaluate the assembly results [26]. MetaGeneMark (Version 3.26) software was used to identify coding regions in the genome using default parameters [27]. Redundancy was removed using MMseqs2 software (Version 12-113e3) using a similarity threshold set to 95% and a coverage threshold set to 90% to construct non-redundant gene catalogue [28, 29].

### 2.6 Taxonomic profiling, functional annotation and microbial data analysis

The protein sequences from the non-redundant gene set were aligned to the NCBI nr database (2019-03) to obtain annotated taxonomic information using Diamond software (version 0.9.24) [30] with a threshold of e-value < 1e-05. The NR that could not be classified to any taxa were defined as unknown taxa. In addition, Eukaryota and Metazoa with very low abundance were excluded. Protein sequences from non-redundant gene catalogue were aligned to the KEGG (Kyoto Encyclopedia of Genes and Genomes) database [31] using Diamond software (version 0.9.24). The threshold was e-value < 1e-05, and if there was more than one match Hit, the best match was selected as the annotation for that sequence. Similarly, the protein sequence of NR was aligned with eggNOG (version4.0) [32] using Diamond software (version 0.9.24). Carbohydrate-activated enzymes (CAZys) were annotated by aligning the protein sequence of NR to the dbCAN database (HMMdb V8) [33] using HMMER (version3.0). The antibiotic resistance genes (ARGs) were annotated by alignment with the Comprehensive Antibiotic Resistance Database (CARD) [34] using RGI (version4.2.2). Protein sequences of NR were annotated by the Virulence Factor Annotation Database (VFDB) [35] using BLAST (version2.2.31+) software [36].

Abundance calculations for sequencing were referred to the previous method [37]. Based on taxonomic, KO and CAZys annotations, the relative abundance of phylum, family, genus, species, KO and CAZys was calculated by summing up the abundance of genes belonging to each category [14]. In addition, the abundance of virulence factor and antibiotic resistance genes in each functional group was calculated from the number of related genes normalized to the number of total NCBI nr-annotated genes in the functional group. Based on the R language (v3.1.1) and python2, alpha diversity analysis was applied via the picante package (v1.8.2); the veganb package (v2.3-0) was applied for the analysis of PERMANOVA/ANOSIM; the pheatmap package (v1.0.2) was applied for the plotting of heat maps. Venn Diagram (v1.6.9) was applied for the analysis of vennd; mothur (v1.22.2) was applied for the analysis of rarefaction curve; matplotlib (v1.5.1) was applied for the analysis of taxonomic composition.

### 2.7 Metagenome-assembled genomes

Metagenomic binning was applied to assembled data by using three different algorithms: MetaBAT252 (version2.12) [38], MaxBin (version2.2.6) [39] and CONOCOCT (version1.0.0) [40]. Subsequently, the software DAS_Tool (version1.1.2 --search_engine diamond --write_bins 1 --score_threshold 0) was used to integrate the results of the different metagenomic binning software [41]. The integrated metagenomic bins (or metagenome-assembled genomes, MAGs) were evaluated using checkM (version1.1.3, default parameters) [42] software. High quality bins were defined by selecting completeness ≥ 80 contamination ≤ 10 (ref Robert D. Stewart) [43].MAGs were clustered into species-level genomic bins (SGBs) using dRep (version3.0.3) [44] with a threshold of 95% ANI. Finally, the GTDB-Tk (v1.2.0) software was used to annotate bins for taxonomic classification by reference to the Genome Taxonomy Database (GTDB) [45]. SGBs containing at least one MAGs in the GTDB were considered to be known SGBs, otherwise, the SGB was considered as unknown [46].

### 2.8 Weighted gene co-expression network analysis

We performed weighted gene co-expression network analysis (WGCNA) using WGCNA package [47] on the platform BMKCloud (www.biocloud.net). Firstly, the expression of all genes in the gut and liver was used to construct gene co-expression modules. Secondly, Pearson correlation correlations between module eigengenes(MEs)and ecological groups were calculated.

### 2.9 Co-occurrence network analysis

The relative abundance of the core microbiota was applied to construct a matrix of correlations, followed by random matrix theory to determine the threshold of correlation. The correlation network data was visualized through the igraph (version 1.2.7) package.

### 2.10 16S rRNA gene sequencing

The composition of the microbiota of largemouth bass was analyzed using 16S rRNA gene amplicons based on the NovaSeq sequencing platform. In brief, the V3-V4 regions of the 16S rRNA genes were amplified by primers [42]. After construction of the libraries, the paired-end 250-nucleotide reads were obtained using the Illumina NovaSeq platform. Using the DADA2 [48] plugin, raw sequences were trimmed, quality filtered, denoised, merged, chimera, and dereplicated. Reads with 97% percent nucleotide sequence identity across all samples were assigned to operational taxonomic units (OTUs) [43]. QIIME2 was applied to the subsequent analysis [49].

### 2.11 Statistical analysis

The Unpaired t test was used to assess the differences between the two groups, and the Tukey’s multiple comparisons test was used to analyze the three or more groups. The plots and diagrams were displayed by ggplot2 (2.2.1) using R language. GraphPad Prism (version 8.0) software was applied for graphing.

## 3. Results

### 3.1 Establishment and assessment of grass carp microbiome gene catalogues

Shotgun metagenomics yielded 476,855,093 valid Reads (Table S1). 2,099,912 contigs were assembled after quality control and de-hosting; 575,856 non-redundant (NR) genes were identified with an average length of 1.65 kb (Table S2). Rarefaction analysis of all samples showed curves close to saturation (Supplementary Fig. 1B). 374,843 NR genes (65.09%) can be blasted to the NCBI-nr database (Table S3). 64.88% of the NR genes could be taxonomically classified. Among them, 97.28% were assigned to bacteria, with the remaining genes being assigned to viruses (1.41%), fungi (1.42%) or archaea (0.17%) (Fig. 1A). At the phylum level, most of the annotated genes (53.74%) belonged to Proteobacteria, followed by Bacteroidetes (16.99%), Firmicutes (13.81%) and Fusobacteria (6.86%) (Fig. 1B). In contrast, Firmicutes and Bacteroidetes are predominant in the human and pig guts, and Proteobacteria makes up a smaller percentage [14], confirming a major compositional difference of fish microbiota compared with mammals. Our further analysis revealed that 48.71% of the NR genes were assigned to bacterial genera, predominantly in *Aeromonas* (17.04%), *Bacteroides* (6.94%), *Shewanella* (5.33%) and *Cetobacterium* (5.1%) (Fig. 1C), which are also different from the case in mammals [50].

**Figure.1.**
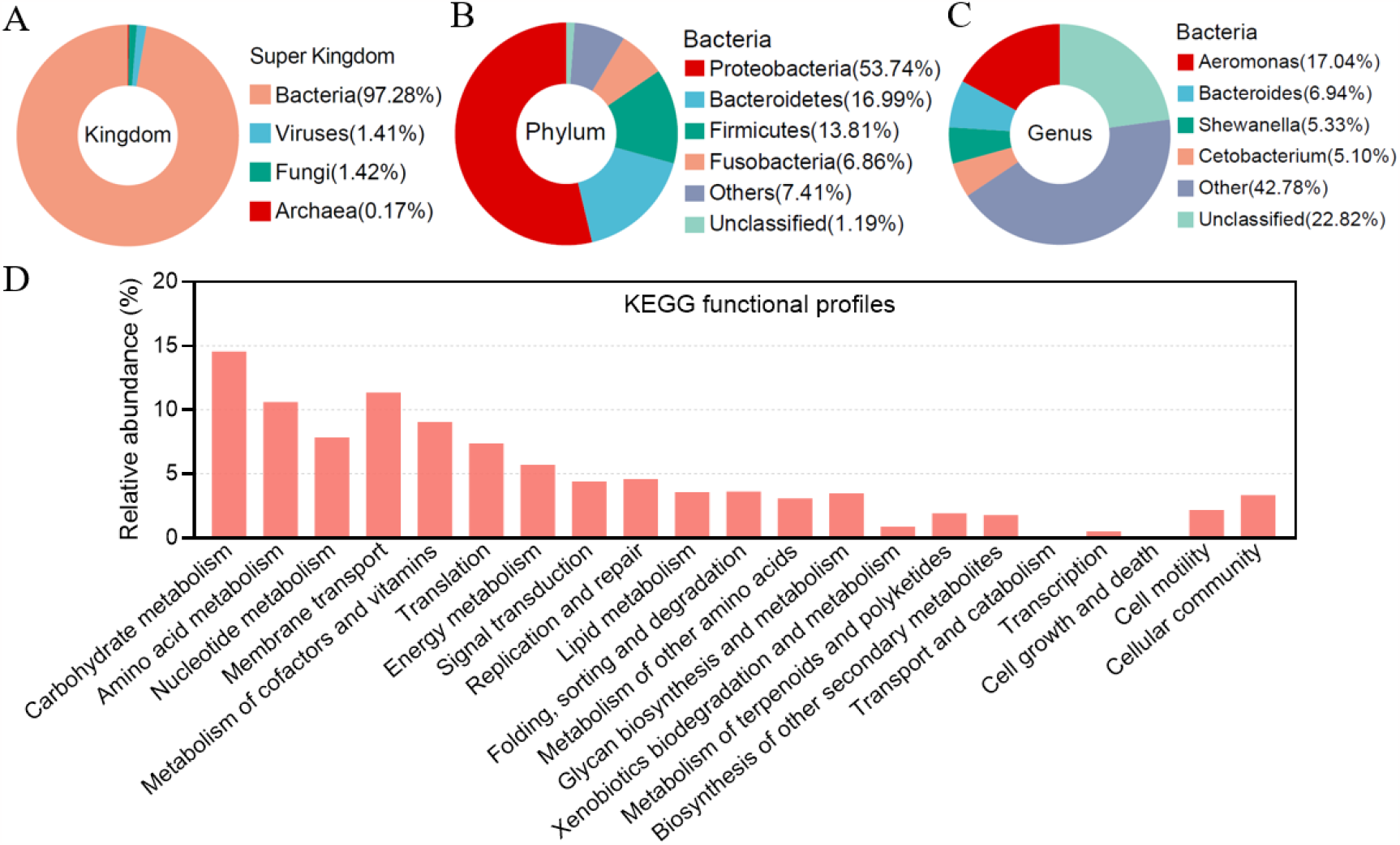
Establishment and analysis of grass carp gut microbiome gene catalogues. (A) Taxonomic annotation of the grass carp gut gene catalog at the super kingdom level. Only non-redundant genes that could be classified from the NCBI nr database were involved in the analysis. (B and C) The NR genes assigned to Bacteria was taxonomically annotated at the phylum (B) and genus (C) level. (D) KEGG functional files of the grass carp gut gene catalogue. Genes without functional annotation were excluded, and relative gene abundance analysis was performed at the second level of the KEGG.

Using KEGG and eggNOG for function classification, 187,972 (32.64%) and 329,336 (57.19%) NR genes were annotated with KEGG orthologous groups (KOs) (Table S4) and eggNOG orthologous groups (OGs) (Table S5). The proportion of genes annotated to KOs or OGs is lower than that in mammals [51], reflecting that gut commensal bacteria were less studied in fish. The KEGG function profiles showed similarities in gut microbial functions of grass carp compared with human and pig. However, the genes for carbohydrate metabolism, amino acid metabolism, nucleotide metabolism and energy metabolism are more abundance in human and pig guts, while the genes for cell motility and cellular community are more abundant in the fish microbiota (Fig. 1D) [14]. These results suggest a deviation of the gut microbial function of fish compared with mammals, with a general lower genetic capacity for nutrient metabolism but higher capacity for microbe-microbe interactions.

Reconstruction of microbial genomes from metagenomic sequencing data has been reported in humans [52], ruminants [43] and pigs [51, 53] by using metagenomic binning methods. We attempted to construct metagenome-assembled genomes (MAGs) from the fish gut based on metagenomic data and assembled a total of 129 high-quality MAGs (Completeness >80% and contamination <10%)(Table S6). The 129 MAGs were further organized into species-level genome bins (SGBs) by average nucleotide identity (ANI) threshold of 95%. This resulted in a total of 18 SGBs. Fifteen SGBs were without any publicly annotatable genomes and were defined as unknown SGBs (uSGB). Subsequent taxonomic annotation by the Genome Taxonomy Database Toolkit (GTDB-Tk) revealed that the 18 SGBs are mainly from Proteobacteria, Fusobacteria, Bacteroidetes and Firmicutes (Table S7).

### 3.2 The ecological interactions of gut microbiota of grass carp

The top 30 genera contributed more than 80% to the total abundance of the microbiota (Supplementary Fig. 2A and B), while the VEEN analysis showed that 23 genera were shared among different dietary groups and time points (Supplementary Fig. 2C). Further analysis among the taxonomically annotated genera revealed that shared genera contributed 79.4%-89.20% of the total abundance (Fig. 2A), indicating that shared genera dominated the microbial community and thus can be considered as core genera. Co-occurrence network analysis was performed to further explore the ecological interactions between the core genera. At week 1, co-exclusive patterns were observed between the core genera belonging to Proteobacteria (except *Acinetobacter*) and Fusobacteria, Bacteroidetes or Firmicutes, while the genera belonging to the same phylum tended to show positive correlations with one another (Fig. 2B). At week 3, co-occurrence network analysis revealed that the genera of Proteobacteria (except *Photobacterium*) had exclusive relations with one or more of the genera of Bacteroidetes, Firmicutes (except *Enterococcus*) or Fusobacteria (Fig. 2C). Similarly, analysis at the phylum level showed that there were mutually exclusive correlations between Proteobacteria and Fusobacteria, Firmicutes or Bacteroidetes (Fig. 2D and E). Thus, the results indicated mutually exclusive relations between the core genera belonging to Proteobacteria and those belonging to Fusobacteria, Firmicutes and Bacteroidetes, suggesting two independent ecological groups of the intestinal microbiota of grass carp: Proteobacteria and Fusobacteria/Firmicutes/Bacteroidetes.

**Figure.2.**
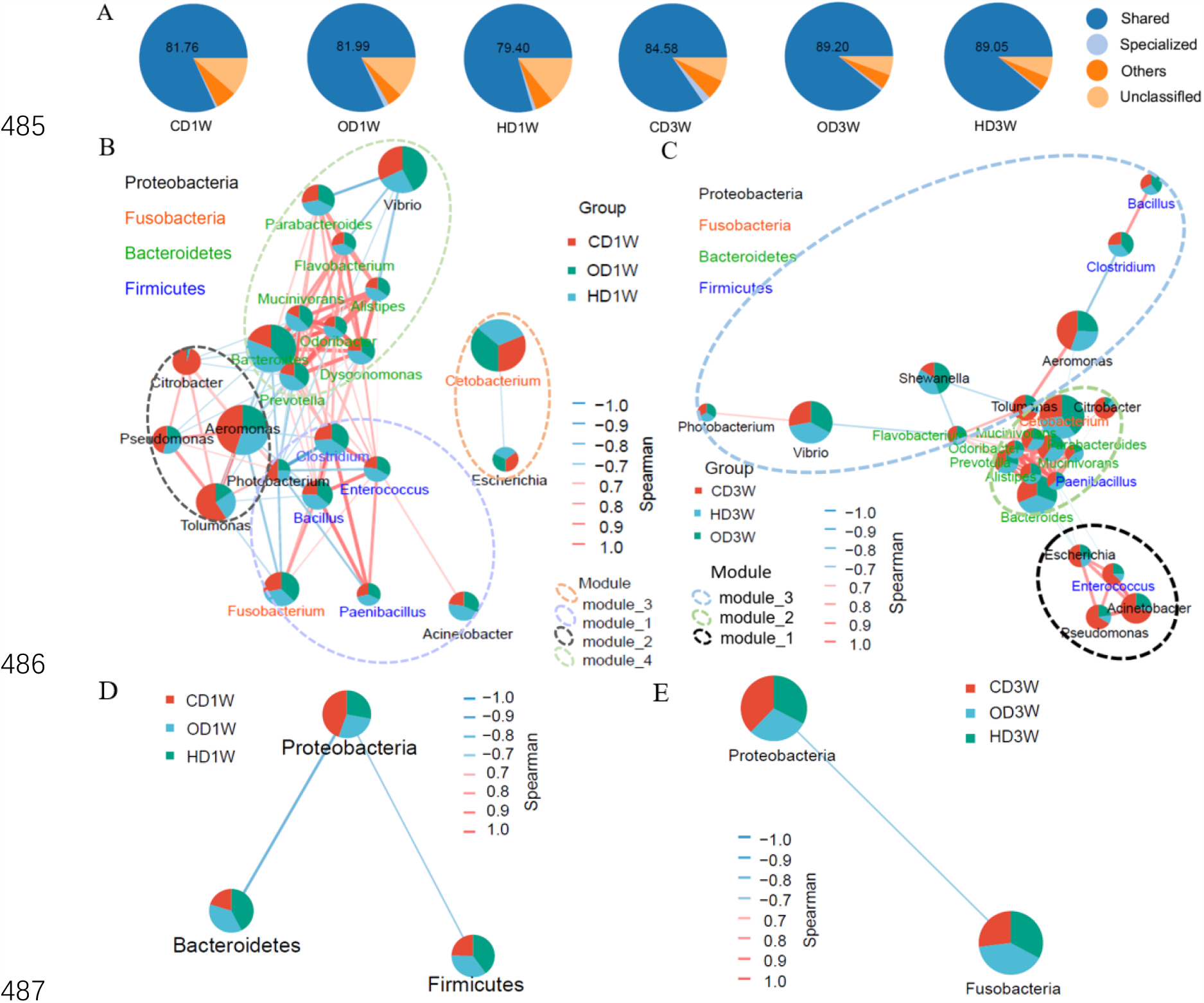
Co-occurrence network analysis of the core genera of grass carp gut microbiota. (A) The relative abundance of shared genera in the microbiota. (B and C) Co-occurrence network of core genera at week 1 and 3. (D and E) Co-occurrence network of microbiota at the phylum level at week 1 and 3. The connection strength threshold is Spearman’s r > 0.5 and correlation is considered as significant when *p*-value < 0.05. The size of the connection point represents the relative abundance of a specific microbe. Red lines indicate co-occurrence. Green lines indicate co-exclusion, and the thickness of the line shows the strength of the correlation. The dotted ellipse indicates a specific modular unit.

### 3.3 Gut microbiota was associated with host gene modules

In order to explore the functional characteristics of the gut microbiota, correlation analysis of the gut microbiota and host gene modules was conducted. WGCNA analysis revealed 24 and 31 gene modules of gut expressed genes at week 1 and 3, respectively (Supplementary Fig. 2D and F). In the liver, 31 and 25 gene modules were obtained at week 1 and 3, respectively (Supplementary Fig. 2E and G).

The association pattern of Proteobacteria with the gene modules of gut and liver at week 1 were consistently opposite to that of Fusobacteria, Firmicutes and Bacteroidetes (Fig. 3A and B), and the opposite association was also observed at week 3 (Fig. 3C and D), indicating differential functionality of Proteobacteria and Fusobacteria/Firmicutes/Bacteroidetes in terms of their interaction with fish host. Therefore, the ecological groups can be considered as two functional groups, i.e., Functional Group 1: Proteobacteria; Functional Group 2: Fusobacteria/Firmicutes/Bacteroidetes. The gut gene modules associated with the microbiota mainly included MEblack, MEgreenyellow, MEpink, MEblue, MEdarkred (week 1) and MEgreen, MEwhite, MEturquoise, MEcyan, MEblack (week 3), while in the liver the main associated gene modules were MElightyellow, MEgreenyellow, MEpurple, MEbrown, MEred, MEmagenta, MEturquoise, MEgreen, MEpink, MElightcyan, MEdarkturquoise (week 1) and MEgreenyellow, MEmagenta, MEroyalblue, MEblue, MEpurple, MEdarkred (week 3) (Fig. 3E).

**Figure.3.**
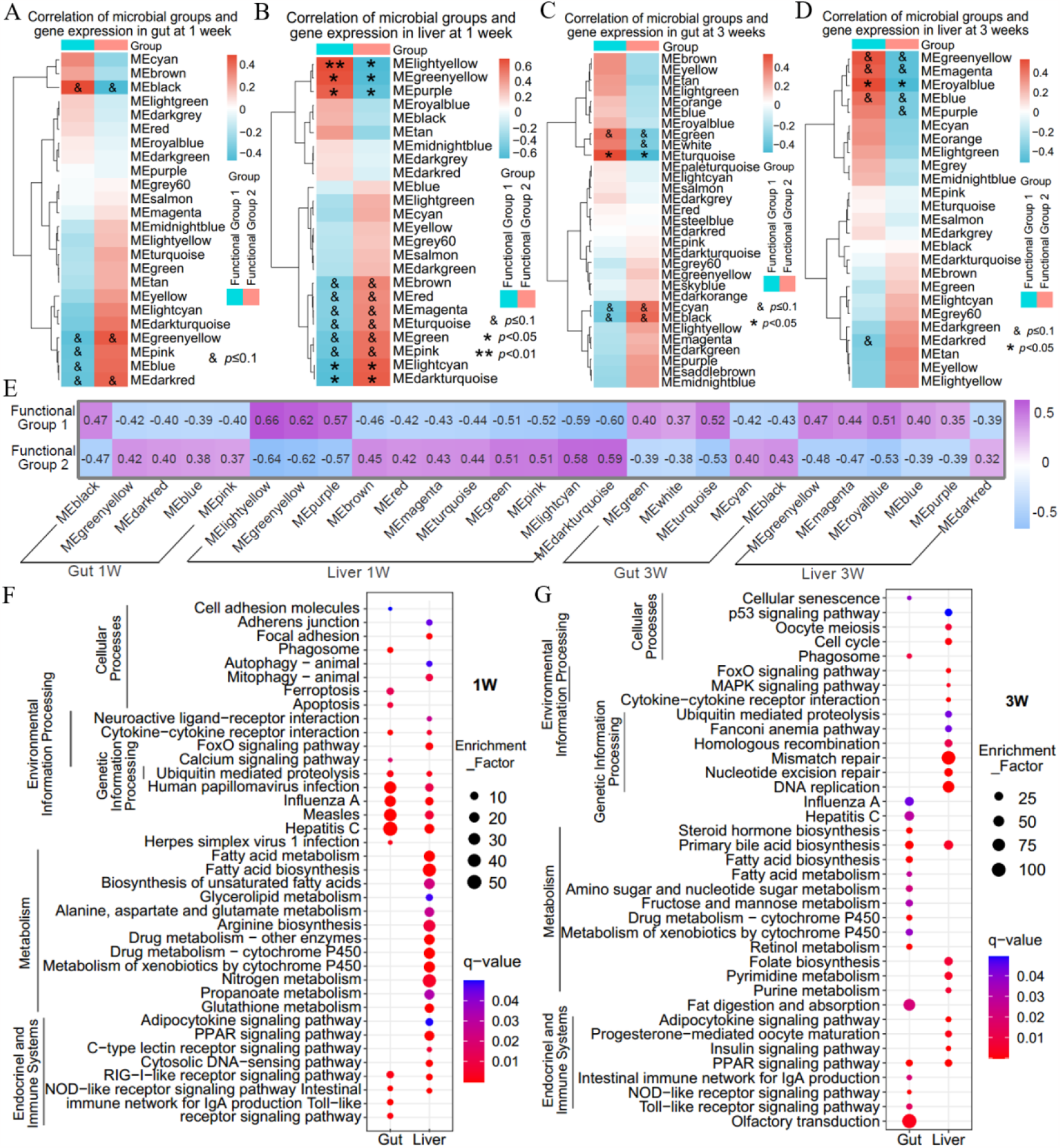
Gut microbiota is associated with host gene modules. (A, B, Cand D) Weighted gene co-expression network analyses (WGCNA) were performed to identify co-expressed gene modules, and the gene modules were correlated with the two ecological groups of the microbiota. A and B represent gut and liver gene modules at week 1, and C and D represent gut and liver gene modules at week 3. Heatmaps show the correlation between gene modules and microbial functional groups. Clustering was performed by a complete clustering method using Euclidean distances. (E) Gene modules with significant correlation with the microbial functional groups. The color of the blocks reflects the value of the correlation coefficient. (F and G) KEGG pathways enriched in the gene modules correlated with the functional groups. Enriched bubble plots have been shown for pathways with q-values < 0.05. Five or six biological replicates were included during analysis, $ *p*< 0.1; * *p*< 0.05; ** *p*<0 .01; *** *p* <0 .001.

KEGG analysis revealed that gut gene modules significantly associated with the microbiota mainly included functions of recognition of microorganisms by the immune system (e.g., Toll-like receptor signaling pathway, NOD-like receptor signaling pathway and RIG-I-like receptor signaling pathway) and their downstream cellular processes (e.g., apoptosis, phagosome, cell adhesion molecules and ferroptosis, etc.) at week 1 (Fig. 3F). In the liver, gene modules with significant association with microbiota involved functions of nutrient metabolism (e.g., propanoate metabolism, fatty acid metabolism, fatty acid biosynthesis and biosynthesis of unsaturated fatty acids, etc.), the immune system (e.g., NOD-like receptor signaling pathway and RIG-I-like receptor signaling pathway, etc.), as well as the endocrine systems (e.g., PPAR signaling pathway and adipocytokine signaling pathway) (Fig. 3F). Similarly, the gut gene modules included functions of the immune system (e.g., Toll-like receptor signaling pathway, NOD-like receptor signaling pathway and RIG -I-like receptor signaling pathway), nutrient metabolism pathways (e.g., fatty acid biosynthesis, fatty acid metabolism and fat digestion and absorption, etc.), and the endocrine system pathway (PPAR signaling pathway) at week3 (Fig. 3G), while in liver the gene modules involved nutritional metabolic pathways (e.g., purine metabolism, pyrimidine metabolism and primary bile acid biosynthesis) and endocrine systems involved in metabolism (insulin signaling pathway, PPAR signaling pathway and adipocytokine signaling pathway) (Fig. 3G). Taken together, the two functional groups were associated with host nutrient metabolism and immunity.

### 3.4 The two functional groups differ in genetic capacity for carbohydrate utilization, virulence factors and antibiotic resistance

The gut microbiota utilizes carbohydrates by carbohydrate-active enzymes (CAZy) and produces SCFAs, which are beneficial to the host in both nutritional and immune aspects [14, 54, 55]. In contrast, virulence factors and antibiotic resistance are generally negative factors that may exert detrimental effect [56-58]. We therefore analyzed the genes encoding CAZy, virulence factors and antibiotic resistance harbored by the two functional groups. The results showed that members of Functional Group 2 enriched CAZy genes encoding enzymes degrading arabinoxylan, pectin, mucin, inulin, and cellulose compared with Functional Group 1, and only the starch-related CAZy gene family was enriched in Functional Group 1 (Fig. 4A and B).

**Figure.4.**
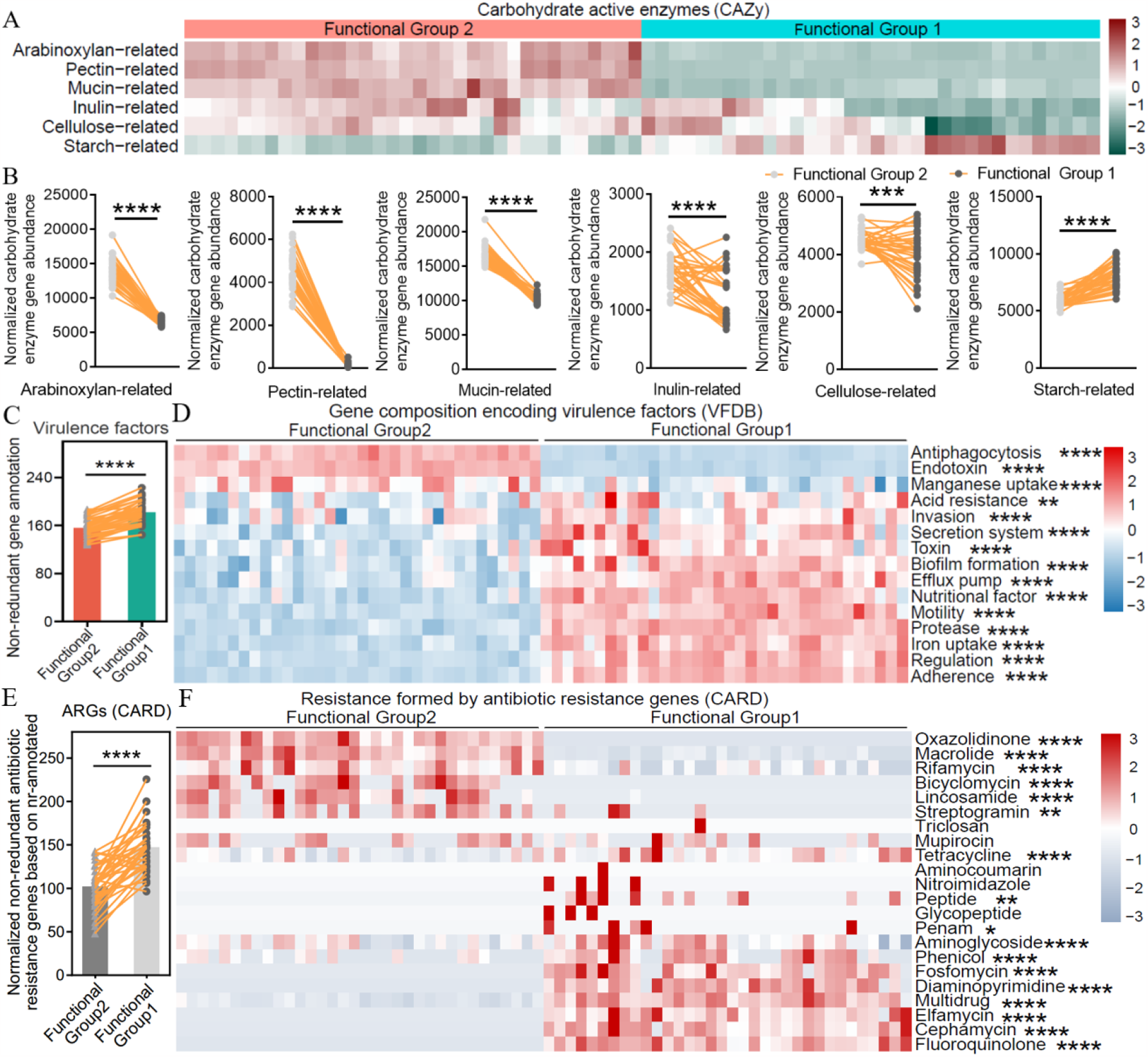
The two functional groups differ in genetic capacity for carbohydrate utilization, virulence factors and antibiotic resistance. (A and B) Differences in genetic capacity for carbohydrate substrate utilization. (A) Heat map showing the proportion of each carbohydrate-active enzyme (CAZy) category in different functional groups. (B) Abundance of genes involved in carbohydrate substrate utilization (The abundance of genes annotated to different types of carbohydrates was normalized to the total carbohydrate-active enzyme in each sample. The abundance of total carbohydrate-active enzyme genes in different functional groups was adjusted to 100,000). Arabinoxylan-related CAZy families: CE1, CE2, CE4, CE6, CE7, GH10, GH11, GH115, GH43, GH51, GH67, GH3 and GH5; pectin-related CAZy families: CE12, CE8, GH28, PL1 and PL9; mucin-related CAZy families: GH1, GH2, GH3, GH4, GH18, GH19, GH20,GH29, GH33, GH38, GH58, GH79, GH84, GH85, GH88, GH89, GH92, GH95, GH98, GH99, GH101, GH105, GH109, GH110,GH113, PL6, PL8, PL12, PL13 and PL21; inulin-related CAZy families: GH32 and GH91; cellulose-related CAZy families: GH1, GH44, GH48, GH8, GH9, GH3 and GH5; starch-related CAZy families: GH13, GH31 and GH97. (C) Number of genes annotated to virulence factors in the two functional groups. (D) The abundance of genes encoding virulence factors in the two functional groups. Genes for each VF class were normalized to the number of total NCBI nr-annotated genes in each functional group. (E) The number of antibiotic resistance genes in the two functional groups. Antibiotic resistance genes were normalized to the total NCBI nr-annotated genes in each functional group (adjusted to 100,000). (F) The abundance of ARGs in the two functional groups. Genes for each resistance class were normalized to the number of total NCBI nr-annotated genes in each functional group. Thirty four biological replicates were included in each functional group during analysis. The Mann-Whitney test was used to analyze differences between functional groups, * *p*< 0.05; ** *p*<0 .01; *** *p*<0 .001; **** *p*<0 .0001.

Furthermore, members of Functional Group 1 encoded more virulence factor (VF) genes compared with Functional Group 2 (Fig. 4C). In particular, VF genes involved in antiphagocytosis, endotoxin and manganese are enriched in Functional Group 2, while Functional Group 1 enriched VF genes across 12 different VF classes, including acid resistance, invasion, secretion system, toxin, biofilm formation, efflux pump, nutritional factor, motility, protease, iron uptake, regulation and adherence (Fig. 4D), suggesting that Functional Group 1 may exert more negative phynotypes in the host-microbes interaction. In terms of antibiotic resistance genes (ARG), the overall number of ARGs is higher in Functional Group 1 versus Functional Group 2 (Fig. 4E). Functional Group encoded more resistance genes against oxazolidinone, macrolide, rifamycin, bicyclomycin, lincosamide and streptogramin, while Functional Group 1 enriched ARGs for resistance to 11 antibiotic classes, i.e., tetracycline, mupirocin, peptide, penam, aminoglycoside, phenicol, fosfomycin, diaminopyrimidine, multidrug, elfamycin, cephamycin and fluoroquinolone (Fig. 4F).

### 3.5 The ratio of “Functional Group 2/Functional Group 1” reflects the structural and functional characteristics of the microbiota

Dietary ingredients are key factors influencing the gut microbiota [59]. We evaluated the effect of diets on the microbial composition of grass carp. Rarefaction analysis of the samples indicated saturation of sequencing depth (Supplementary Fig. 3A and C). At week 1, the relative abundance of Proteobacteria decreased in OD and HD groups compared with CD group, while the abundance of Firmicutes and Bacteroidetes increased (Fig. 5A). At the genus level, the abundance of *Aeromonas* and *Vibrio* decreased while *Cetobacterium* and *Bacteroides* increased in OD and HD groups versus CD group (Supplementary Fig. 3B). Similarly, the abundance of Proteobacteria deceased and Fusobacteria increased in OD and HD groups compared with CD group at week 3 (Fig. 5B), with an increase in the abundance of *Cetobacterium* and decease of *Aeromonas* in OD and HD groups versus CD (Supplementary Fig. 3D). Diets had no significant influence on the α-diversity index of the microbiota (Supplementary Fig. 3E and F). β-diversity analysis revealed a significant alteration of the microbiota due to dietary groups at both week 1 and 3. The microbiota of CD clustered alone, while the microbiota of OD and HD clustered together (Supplementary Fig. 3G, H, I, K and L).

**Figure.5.**
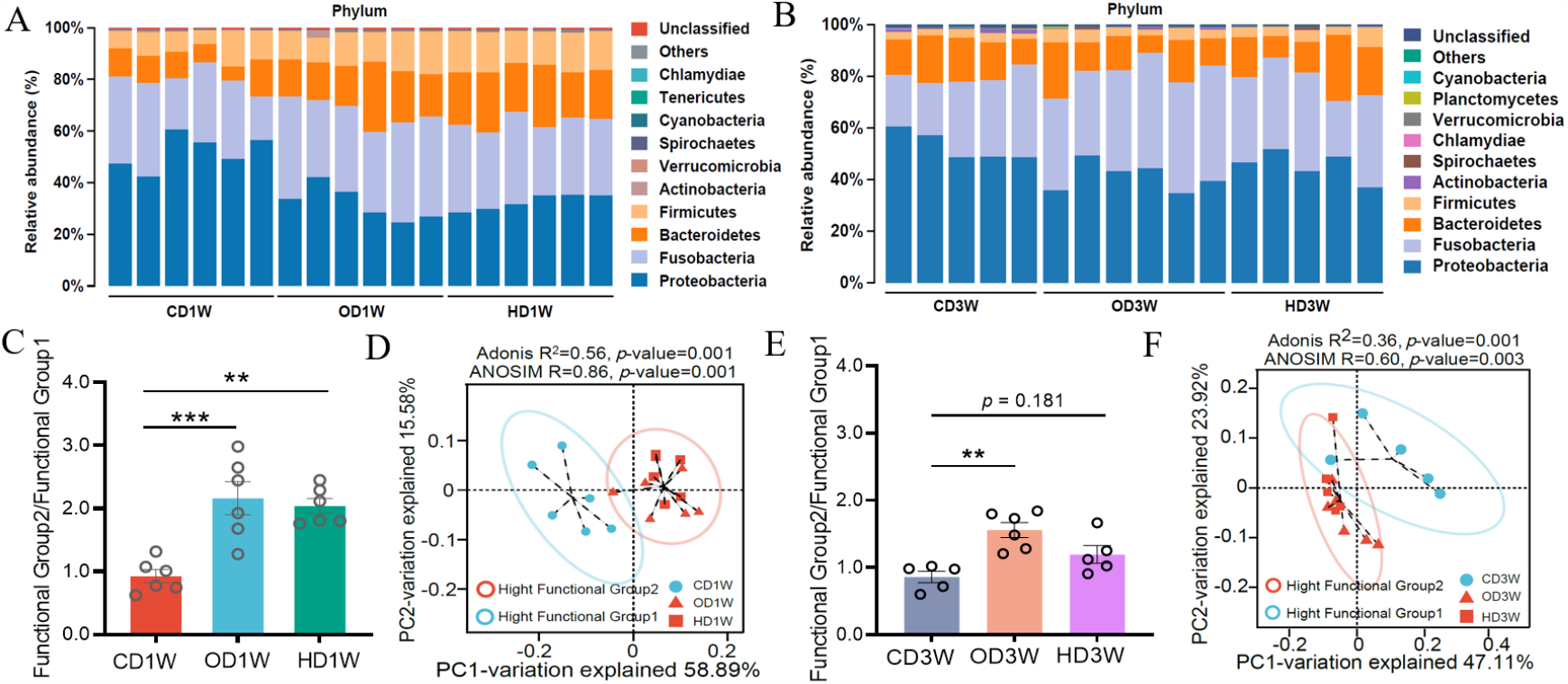
The ratio of “Functional Group 2/Functional Group 1” reflects the structural difference of microbiota associated with different dietary groups. (A and B) The relative abundance of top 10 phylum of the gut microbiota at week 1 and 3. (C and E) Ratio of “Functional Group2/Functional Group1” at week 1 and 3. (D and F) Principal coordinate analysis (PCoA) of the microbiota associated with different dietary groups by Bray-Curtis’s distance. Samples with “High Functional Group2” or “High Functional Group1” were marked with different colors. The dotted ellipse borders represent the 95% confidence interval. Data were expressed as the mean ± SEM (n = 5 or 6 biological replicates), ** *p*<0 .01; *** *p*<0 .001.

Considering the ecological and functional difference of Functional Group 1 and Functional Group 2, the ratio of the abundance of Functional Group 2 and Functional Group 1, designated as “Functional Group 2/Functional Group 1” was calculated to evaluate the structural and functional characteristics of the microbiota. At week 1, “Functional Group 2/Functional Group 1” was significantly higher in OD/HD diets versus CD diet (Fig. 5C), and a similar trend was observed in “Functional Group 2/Functional Group 1” among the three dietary groups at week 3 (Fig. 5E). PCoA analysis showed that the microbial structure associated with different diets was well explained by “Functional Group2/Functional Group 1”, with the microbiota of high ratio (OD/HD) deviating from those of low ratio (CD) at both week 1 (Fig. 5D) and week 3 (Fig. 5F). Thus, the ratio of “Functional Group2/Functional Group 1” can be used as a parameter to evaluate the structural characteristics of the gut microbiota of grass carp, which efficiently reflected the overall effect of diets on the microbial composition.

Furthermore, PCoA analysis showed a robust separation between the microbiota of high ratio groups (OD/HD) and low ratio CD group in terms of the abundance of genes encoding carbohydrate-coding enzymes (ANOSIM, R=0.85, *p*=0.001; R=0.51, *p*=0.007), and the separation was efficient at both week 1 and 3 (Fig. 6A and B).Compared with low ratio microbiota (CD), high ratio microbiota (OD/HD) enriched CAZy genes for arabinoxylan, pectin, and cellulose utilization while the CAZy genes for starch utilization was decreased (Fig. 6C and D). On top of it, the microbiota with high ratio harbored more abundance of the gene encoding the key enzyme (FTHFS) for acetate production. A similar trend was observed for butyrate producing key enzymes (Buk, AtoA/D), while no obvious difference was observed for the propionate producing gene (PcoAt) between the high ratio (OD/HD) and low ratio (CD) groups (Fig. 6E). Together, these results indicated that the microbiota of high ratio groups had higher functional capability for carbohydrate utilization and SCFAs production.

**Figure.6.**
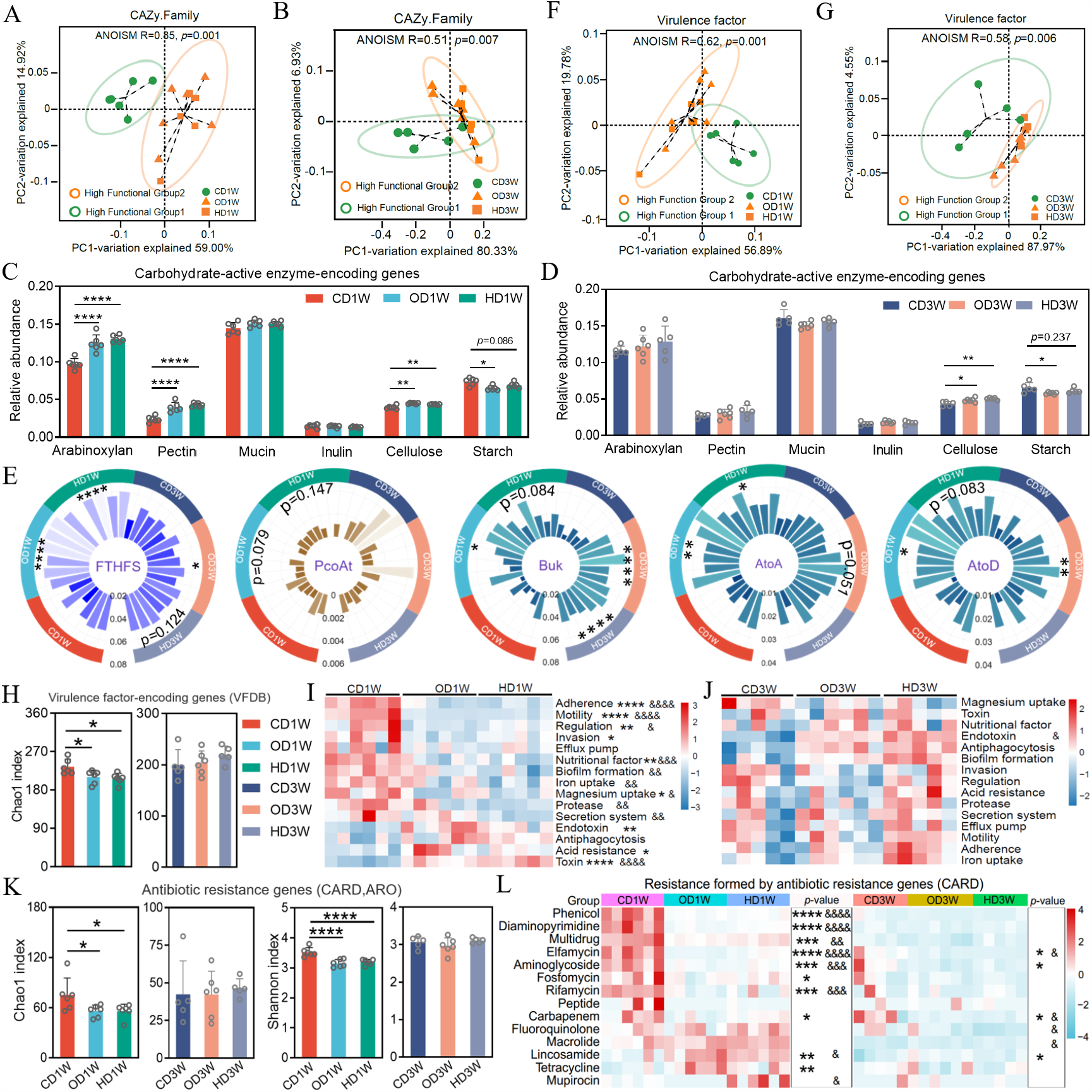
The ratio of “Functional Group 2/Functional Group 1” reflects the functional characteristics of the microbiota. (A and B) PCoA analysis of the abundance of CAZy genes in different dietary groups. Samples with “High Functional Group 2” or “High Functional Group 1” were marked with different colors. (C and D) Relative abundance of CAZy genes for a particular substrate in different dietary groups. (E) The relative abundance of key enzyme genes associated with the production of acetate (FTHFS: acetyl-formyltetrahydrofolate synthase/formate-tetrahydrofolate ligase), propionate (PcoAt: propionyl-CoA:succinate-CoA transferase/propionate CoA-transferase) and butyrate (Buk: butyrate kinase, AtoA: acetoacetate CoA transferase beta subunit and AtoD: acetoacetate CoA transferase alpha subunit) in different dietary groups based on KEGG orthologous groups. (F and G) PCoA analysis of genes encoding virulence factors in different dietary groups. Samples with “High Functional Group 2” or “High Functional Group 1” were marked with different colors. (H) Chao1 index analysis of virulence factor gene abundance. (I and J) The relative abundance of each class of virulence factor genes in different dietary groups at week 1 (I) and week 3 (J). (K) The richness and diversity of antibiotic resistance genes were analyzed by Chao1 and Shannon indices, respectively. (L) The relative abundance of each class of antibiotic resistance genes in different dietary groups. ARGs with relative abundance below 1 in 100,000 were excluded in the analysis.

The expression of GPCR43 in gut was higher in high Functional Group 2 diets (OD/HD) compared with CD (Supplementary Fig. 4A). In the gut, acetyl-CoA production from pyruvate, amino acids (leucine, isoleucine) and fatty acids was all suppressed in dietary groups with high Functional Group 2 (OD/HD) versus high Functional Group 1 group (CD) (Supplementary Fig. 4B). On the other hand, the expression of ACSS1_2, the key enzyme for acetate assimilation, was up-regulated. Similarly, high Functional Group 2 groups (OD/HD) featured reduced acetyl-CoA production from pyruvate, amino acids (leucine, valine, isoleucine) and fatty acids in liver when compared with high Functional Group 1 group (CD), and the expression of ACSS1_2 was enhanced (Supplementary Fig. 4A). In both gut and liver (Supplementary Fig. 4A and B), the fatty acids synthesis from acetyl-CoA was enhanced in high Functional Group 2 dietary groups (OD/HD) versus CD group, indicating that the exogenous acetate influenced lipid metabolism. Together, these results indicate that the differential SCFAs production capability of the microbiota with different “Functional Group 2/Functional Group 1” ratio affected host metabolism, and acetate was the main SCFA that exert the influence.

Further analysis of the virulence factors revealed significant structural differences in the genes encoding the virulence factors for dietary groups with significant differences in the ratio of “Functional Group2/Functional Group 1” (ANOSIM, R=0.62, *p*=0.001; R=0.58, *p*=0.006) (Fig.6F and G). High ratio groups encoded lower abundance of virulence factor genes compared with low ratio group (Fig.6 H, I and J). This trend was more obvious at week 1 compared with week 3, which accorded with the reduced difference in the ratio among groups. Regarding antibiotic resistance, high ratio groups (OD/HD) had lower abundance and diversity of ARGs compared with low ratio group (CD) (Fig. 6K and L)

### 3.6 The diversity and composition of the gut microbiota were associated with the diversity and composition of DNA viruses

The non-redundant gene catalogue was annotated through the Nr database for the DNA virome, and Rarefaction Curves indicated that the Shotgun metagenomics sequencing depth was adequate to capture the DNA viruses (Supplementary Figs. 5A and B). The analysis revealed that the intestinal DNA virome was dominated by Caudovirales, which consisted of Myoviridae, Siphoviridae, and Podoviridae (Supplementary Figs. 5C and D, Table S12). Compared to the CD group, the relative abundance of Myoviridae increased and Siphoviridae decreased in the OD/HD group at both week 1 and 3 (Supplementary Figs. 5C and D). At the genus level, the gut DNA virome mainly consisted of *Spn3virus, Vhmlvirus, Phikzvirus, Eah2virus*, and *Machinavirus* (Table S12). Compared to CD, the relative abundance of *Spn3virus, Vhmlvirus, Machinavirus*, and *Eah2virus* increased and *Phikzvirus* decreased in the OD/HD dietary groups (Supplementary Figs. 5E and F). Interestingly, the Chao1 and Shannon indexes of the microbiota were strongly positive correlated with DNA virome (Figs. 7A and B; Supplementary Figs. 6A, B and C). A correlation study of beta diversity showed a strong positive correlation between the structural characteristics of the gut microbial community and DNA viruses (Fig. 7C and D), which was stable at both week 1 and 3 (Supplementary Fig. 6D). More specifically, *Aeromonas, Tolumonas* showed negative correlations with phage, while *Bacteroides, Fusobacterium, Clostridium*, and *Bacillus* exhibited positive correlations at week 1, including Spn3virus, Vhmlvirus, Phikzvirus, Machinavirus (Fig. 7E). At week 3, *Vibrio, Aeromonas, shewanella, Acinetobacter* and *Tolumonas* were found to correlate with phages (Fig. 7F). Therefore, the diversity and structure of gut microbiota were significantly correlated with DNA viruses (mainly phages) in grass carp. Similar results were reported in human [60], indicating a conserved correlation of commensal bacteria and phages.

**Figure.7.**
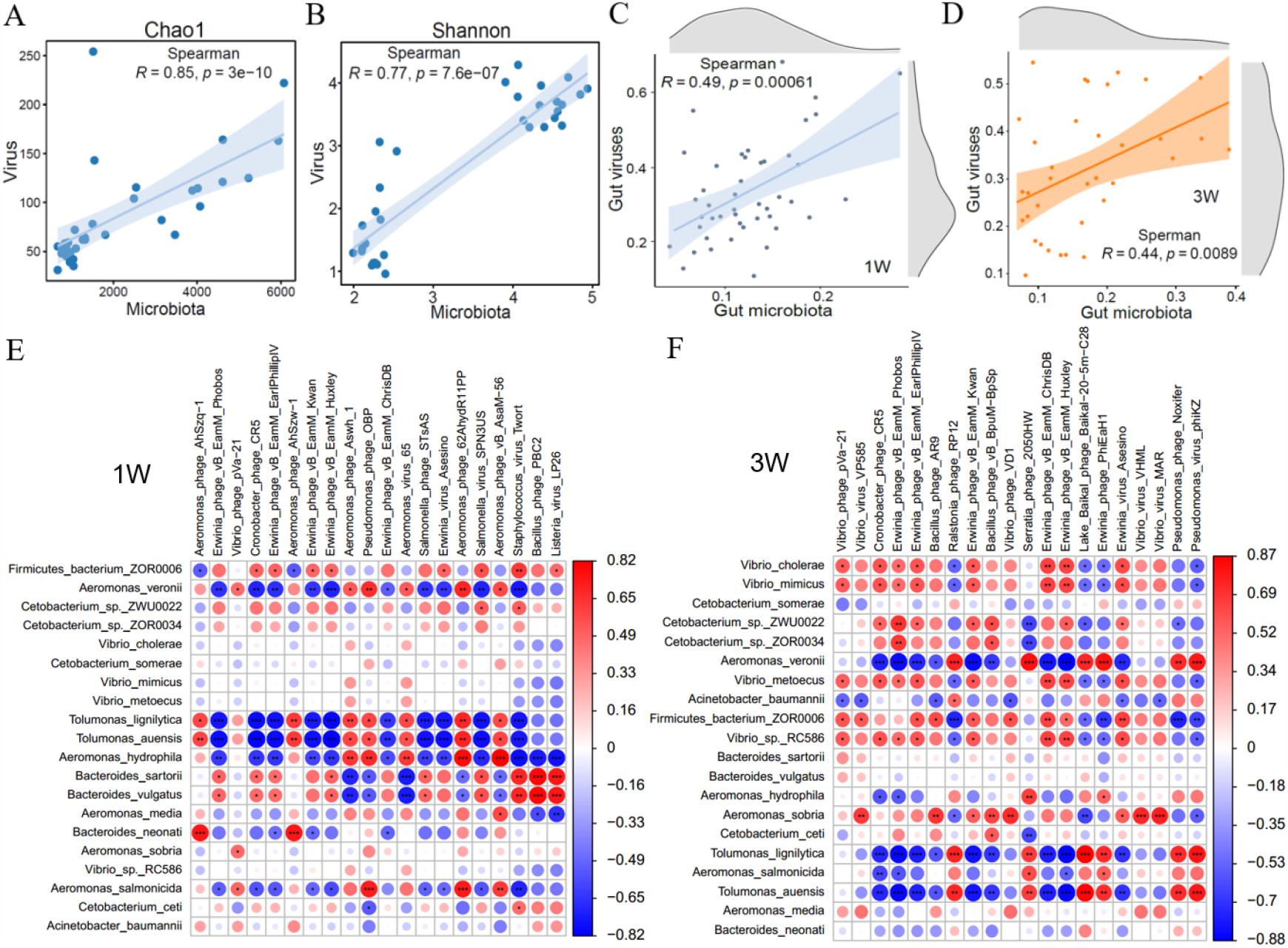
Diversity and composition of DNA viruses were associated with the diversity and composition of the gut microbiota. (A and B) Spearman correlation analysis between gut DNA viral and microbiota alpha diversity at week 1 and 3. (C and D) Correlation analysis of beta diversity of intestinal DNA viruses and microbiota (E and F) The heat map shows the correlation between intestinal DNA viruses (mainly phages) and microbiota at the species level. Correlation was conducted by Spearman’s analysis. Red represents positive correlation; blue represents negative correlation and the shade of the color indicates the value of the correlation coefficient.

## 4. Discussion

The gene catalogue of grass carp reported in this study comprises 575,856 NR genes, which can serve as a reference for further metagenomic studies of this important economic fish species. Moreover, the gene catalogue can be regarded as a milestone of fish metagenomics-related studies, as this is the first fish gut microbial gene catalogue to our knowledge. Despite large compositional difference, functional analysis revealed shared functions of fish microbiota with those of mammals, indicating functional redundancy of the microbiota. Nevertheless, compared to mammals, the microbiota of grass carp featured lower abundance of genes related to nutrient metabolism, suggesting lower contribution of the microbiota to host metabolism. It will be interesting to evaluate whether this is a common feature of fish microbiota, which awaits gut metagenomic analysis of more fish species.

Firmicutes and Bacteroidetes are the dominant phyla in the gut microbiota of human and other mammals [61-63]. In contrast, fish microbiota comprises high abundance of Proteobacteria [64], which is the case in this study for grass carp. Proteobacteria is known to include many pathogenic or opportunistically pathogenic genera/species, and is generally regarded as negative components in the commensal microbiota [65]. However, it is a paradox that this “negative” phylum often occupies high abundance in fish microbiota, and the functionality of Proteobacteria in fish has never been assessed in a systematic way. The results in our study indicate that the functional implication of Proteobacteria is generally negative, as they harbor less genes for carbohydrate degradation and SCFAs production, while encoding more virulence factor and antibiotic resistance genes.

We constructed the co-occurrence network of the core genera and observed that except for a few genera, members from Proteobacteria showed negative correlations with the genera belonging to Firmicutes, Fusobacteria and Bacteroidetes. Therefore, Proteobacteria and Firmicutes/Fusobacteria/Bacteroidetes form two ecological groups. Further studies revealed consistent opposite association pattern of the two ecological groups with gene modules of host, which prompted us to define them as two functional groups. Consistent with their opposite association patters with the host, the two functional groups showed difference in genetic capacity for carbohydrate utilization, SCFAs production, virulence factors and antibiotic resistance. Considering their ecological and functional difference, we proposed that the ratio of “Functional Group 2/Functional Group 1” can be used to evaluate the structural and functional characteristics of grass carp gut microbiota. We found that the ratio was robust in reflecting overall compositional difference of the microbiota associated with different diets. Moreover, the functionality of the microbiota can also be reflected by the ratio. We also found that the ratio of “Functional Group 2/Functional Group 1” can also depict structural characteristics of gut microbiota of other fish species, including zebrafish and largemouth bass (Supplementary Figs. 7 and 8), suggesting potentially extensive application of the ratio in evaluating the gut microbiota of fish species. In mammals, the Firmicutes/Bacteroidetes ratio of intestinal microbiota reflects the degree of obesity [66]. The ratio of “Functional Group 2/Functional Group 1” proposed in this study is a similar way of description of the microbiota. Our preliminary data suggest that the ratio reflects the overall health of fish, with higher ratio correlating with better health (Supplementary Fig. 9 and data not shown), but the specific functional implications of the ratio deserve further detailed studies. Notably, the two functional groups comprise major phyla of the gut microbiome of grass carp, and thus the ratio is a relatively rough biomarker that depicts the characteristics of the microbiota. Further studies are warranted to investigate the ecological and functional interactions of the core sub-groups within the two functional groups, which may give rise to more accurate microbiome signature as reported in human [58].

Enterotypes was first reported to exist in humans with robust clustering of gut microbial communities [61, 67]. Enterotype-like structures have also been reported in several animal studies, although their gut microbial composition differs from that of humans. In addition, there has been found to be a correlation between enterotype and host growth traits in pigs [62]. It should be noted that the microbiota variation of grass carp observed in this study is more like to be continuous, which is not a stratified variation that can be depicted by enterotypes. This might be due to that the influence of “artificial diets” on the intestinal microbiota of grass carp is too big, which overwhelmed the possible stratified variation of the microbiota in natural cases. The ratio of “Functional Group 2/Functional Group 1” can reflect the structural and functional features of fish microbiota, and the microbiota associated with different diets and time points can be roughly classified as “high ratio” and “low ratio”. However, such classification does not fit the concept of enterotypes. Enterotypes that can robustly cluster the gut microbiota variation in fish deserves further investigation.

## 5. Conclusions

The gut microbial gene catalogue of grass carp extended our knowledge about the gut microbiome of this economically important fish species, and provided resources for fish gut microbiome-related research. The taxonomic and functional annotation results reflected that the gut commensal bacteria were less studied in fish compared with mammals, highlighting the importance of more fundamental research in this field. We found that Proteobacteria is generally negatively correlated with the members of Fusobacteria/Firmicutes/Bacteroidetes, and the two groups showed differential functionality in terms of their interaction with fish host. Consistent with their differential functionality, the two functional groups differed in the genetic capacity for carbohydrate utilization, SCFAs production, virulence factors and antibiotic resistance. Furthermore, the ratio of “Functional Group 2/Functional Group 1” can efficiently reflect the structural and functional characteristics of the gut microbiome of grass carp, and could be used as a biomarker to assess the microbiota. Our results provide insights into functional implications of the main phyla that comprise the fish microbiota, and shed lights on targets for microbiota regulation, which may promote the development of green inputs for aquaculture that derive from or target the gut microbiota.

## 7. Ethics approval

All experimental procedures were performed in accordance with the “Guidelines for Experimental Animals” of the Ministry of Science and Technology (Beijing, China). The study was reviewed and approved by the Feed Research Institute of the Chinese Academy of Agricultural Sciences Animal Care Committee under the auspices of the China Council for Animal Care. All efforts were made to minimize suffering.

