## Supplementary data for "Deciphering the gut microbiome of grass carp through multi-omics approach"

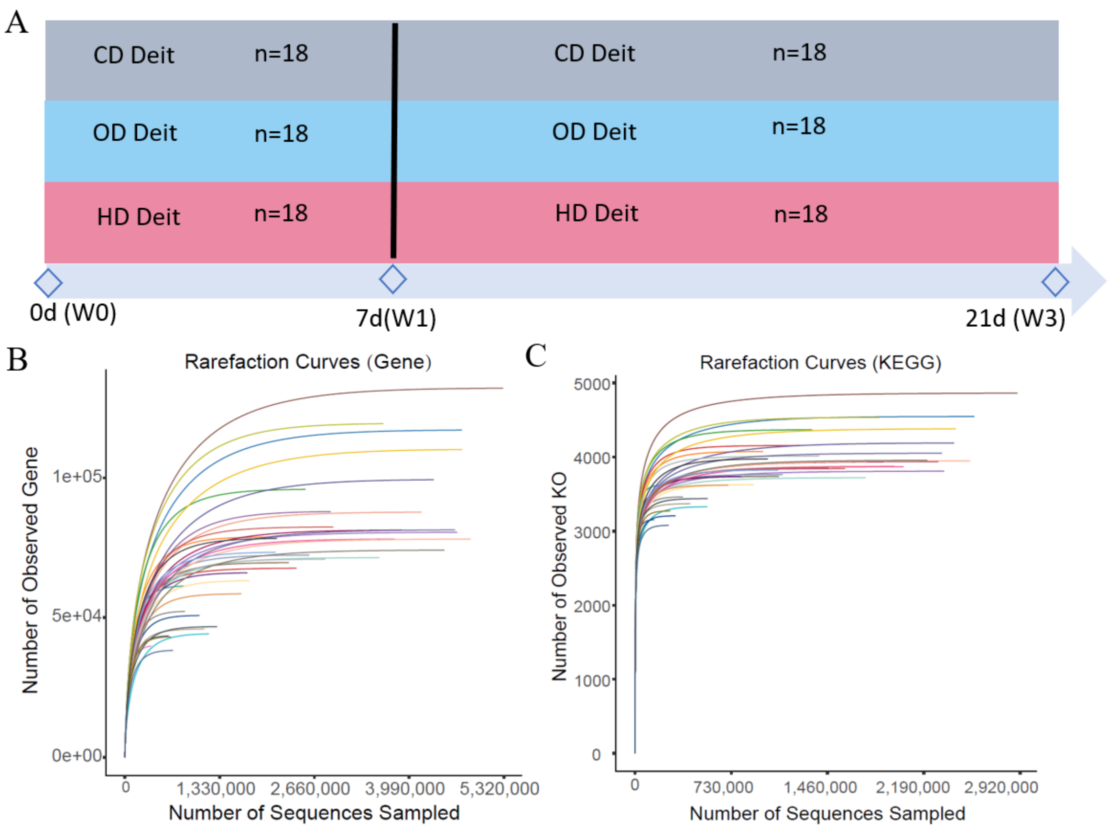


**Figure S.1 Experimental design and rarefaction curve analysis of microbial non-redundant genes.** (A) Experimental design. W1 sampling was 18 fish randomly removed from each group out of 36 and the rest were cultured to three weeks. (B and C) rarefaction curve analysis of gut microbial genes for non-redundant genes and analysis of microbial KO (KEGG orthologous groups) in each sample.

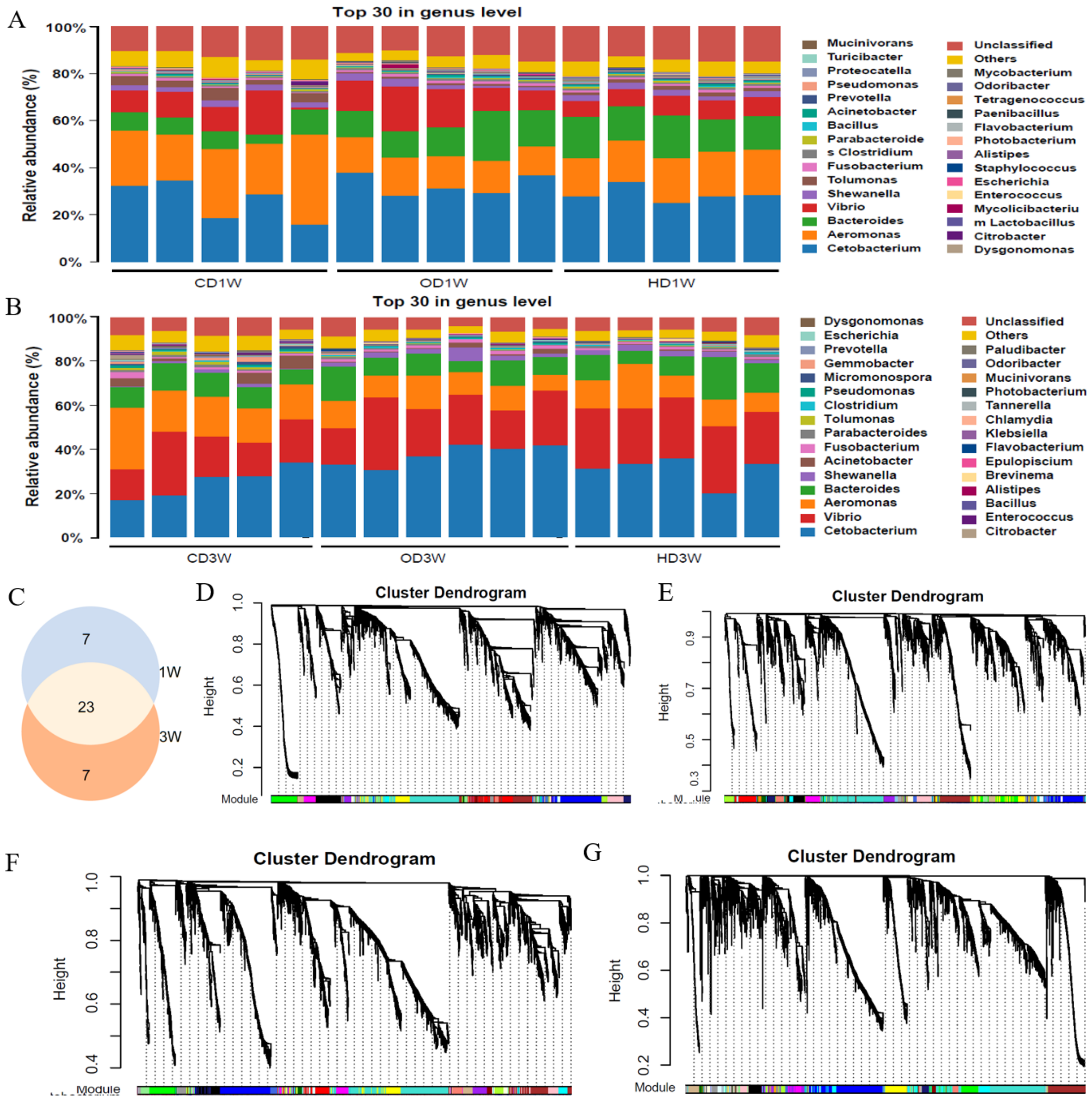


**Figure S.2 WGCNA analysis to reveal the association between host gene expression and gut microbiota.** (A and B) The main composition of the microbiota at the genus level in different dietary groups (n = 5 or 6 biological replicates). (C) The microbial communities shared by different groups at the genus level were analyzed by Venn. WGCNA was performed to explore the gene expression modules in the gut and liver. (D and E) The gut and liver gene modules at week 1 (F and G) The gut and liver gene modules at week 3.

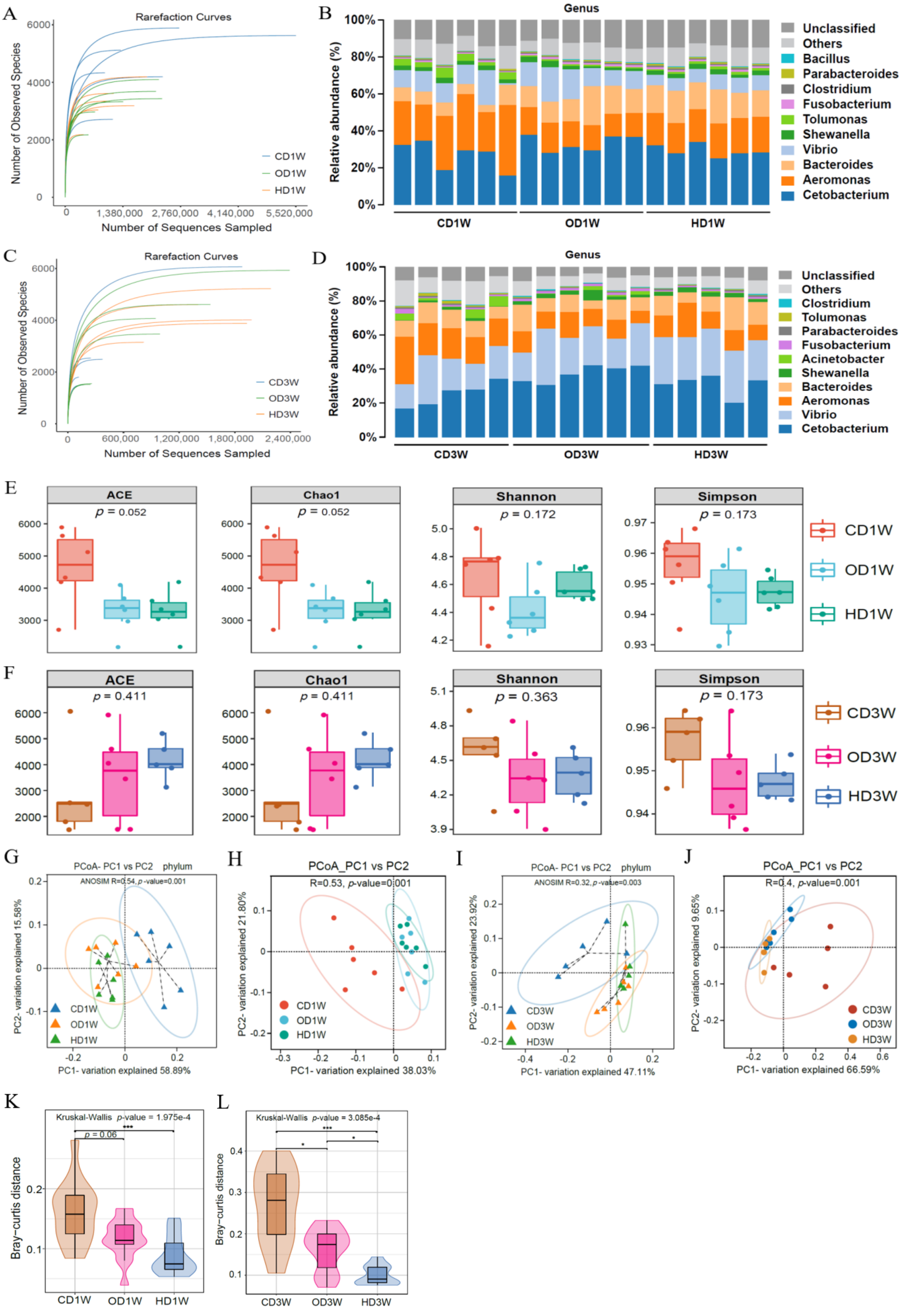

A
Rarefaction Curves
Number of Observed Species
Number of Sequences Sampled
CD1W
OD1W
HD1W
B
Genus
Relative abundance (%)
Unclassified
Others
Bacillus
Parabacteroides
Clostridium
Fusobacterium
Tolumonas
Shewanella
Vibrio
Bacteroides
Aeromonas
Cetobacterium
CD1W
OD1W
HD1W
C
Rarefaction Curves
Number of Observed Species
Number of Sequences Sampled
CD3W
OD3W
HD3W
D
Genus
Relative abundance (%)
Unclassified
Others
Clostridium
Tolumonas
Parabacteroides
Fusobacterium
Acinetobacter
Shewanella
Bacteroides
Aeromonas
Vibrio
Cetobacterium
CD3W
OD3W
HD3W
E
ACE p = 0.052
Chao1 p = 0.052
Shannon p = 0.172
Simpson p = 0.173
CD1W
OD1W
HD1W
F
ACE p = 0.411
Chao1 p = 0.411
Shannon p = 0.363
Simpson p = 0.173
CD3W
OD3W
HD3W
G
PCoA- PC1 vs PC2 phylum
ANOSIM R=0.54, p-value=0.001
PC1- variation explained 58.89%
PC2- variation explained 15.58%
CD1W
OD1W
HD1W
H
PCoA_PC1 vs PC2
R=0.53, p-value=0.001
PC1- variation explained 38.03%
PC2- variation explained 21.80%
CD1W
OD1W
HD1W
I
PCoA- PC1 vs PC2 phylum
ANOSIM R=0.32, p-value=0.003
PC1- variation explained 47.11%
PC2- variation explained 23.92%
CD3W
OD3W
HD3W
J
PCoA_PC1 vs PC2
R=0.4, p-value=0.001
PC1- variation explained 66.59%
PC2- variation explained 9.65%
CD3W
OD3W
HD3W
K
Kruskal-Wallis p-value = 1.975e-4
p = 0.06
***
Bray-curtis distance
CD1W
OD1W
HD1W
L
Kruskal-Wallis p-value = 3.085e-4
***
*
*
Bray-curtis distance
CD3W
OD3W
HD3W

**Figure S.3 Gut microbial composition and diversity of grass carp in different dietary groups.** (A and C) Rarefaction Curves of samples from different dietary groups. (B and D) Relative abundance of the top 10 genera at week 1 and 3. (E and F) Alpha diversity of the gut microbiota at the genus level at week 1 (E) and 3 (F), including ACE, Chao1, Shannon and Simpson indices. (G and I) PCoA analysis of all samples at the phylum level by Bray-Curtis's distance (ANOISM: R=0.54 $p$-value=0.001; R=0.32 $p$-value=0.003 at week 1 and 3, respectively). (H and J) PCoA analysis of all samples at the genus level (ANOISM: R=0.53, $p$-value=0.001; R =0.4, $p$-value=0.003 at week 1 and 3, respectively). The dotted ellipse borders represent the 95% confidence interval. (K and L) Bray-Curtis's distance was calculated based on grouping distance matrix and the result showed that there was a significant difference in microbiota between the OD or HD diets and the CD diet, with less difference between the microbiota of the OD and HD diets. Kruskal-Wallis non-parametric test was conducted to obtain $p$-values for all group differences, followed by Dunn's test for significance levels between the two groups. Data were expressed as the mean ± SEM (n = 5 or 6 biological replicates).

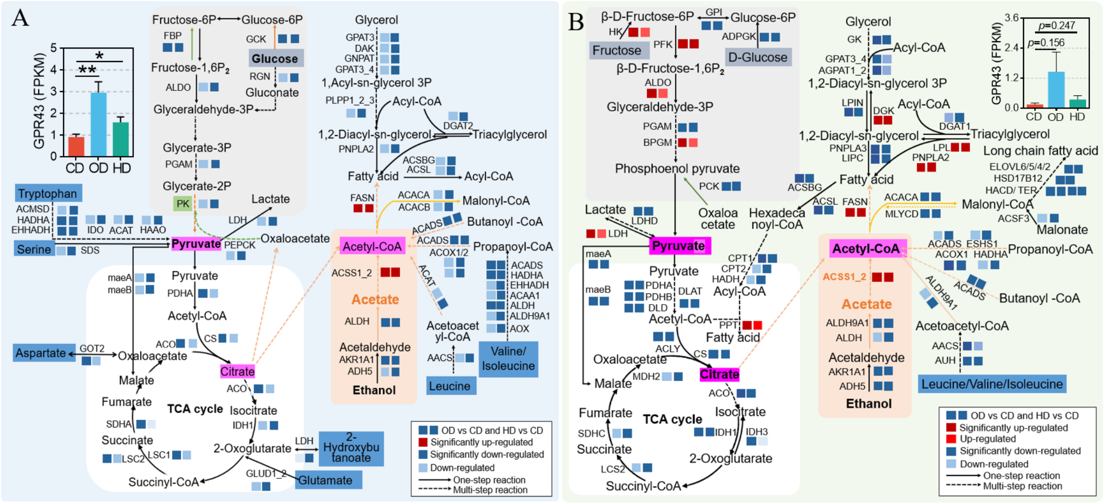


**Figure S.4 Microbiota-derived acetate affected host metabolism.** (A and B) Microbiota-derived acetate affected host metabolism in the gut and liver. The Data were represented with 5 or 6 biological replicates per treatment group (Table S8 and 9).

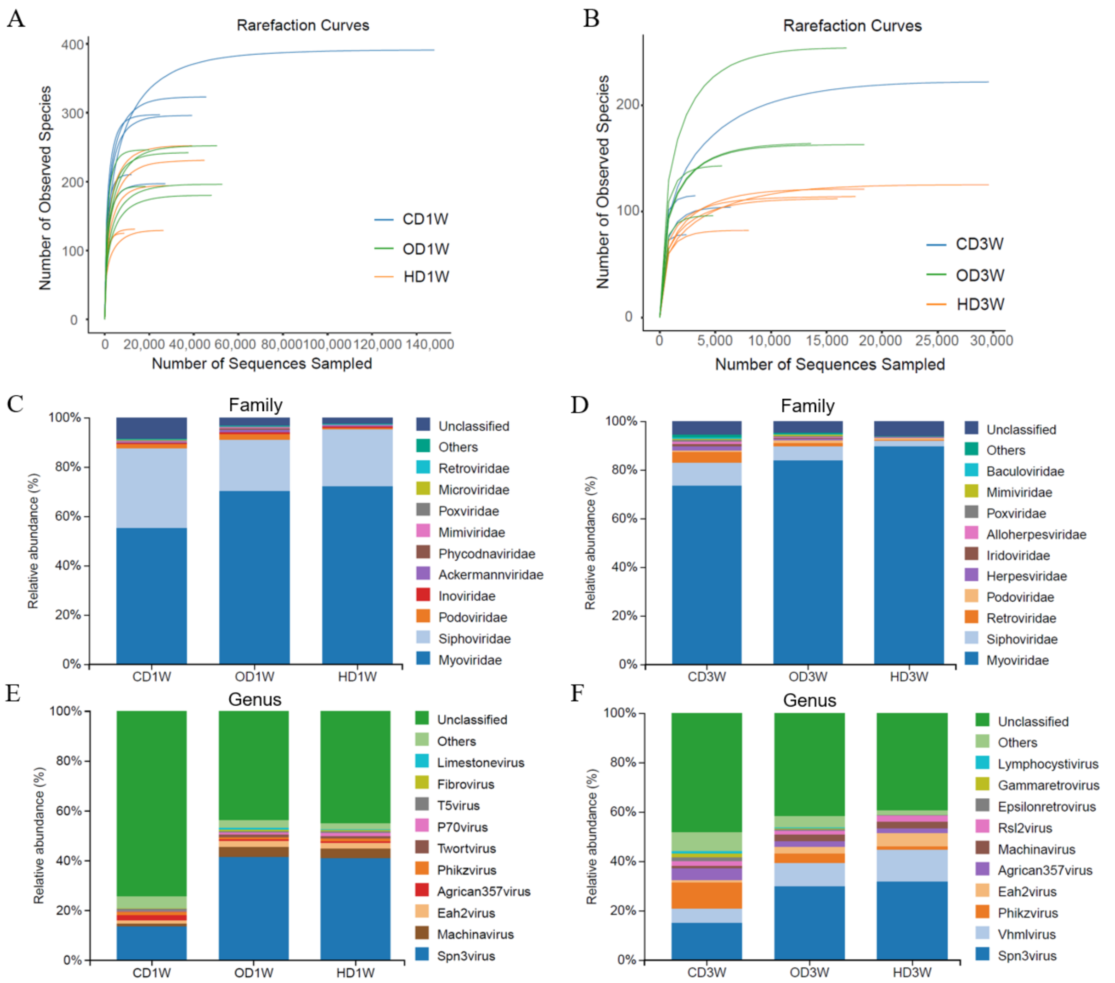


**Figure S.5 Taxonomic composition of intestinal DNA viruses.** (A and B) Rarefaction curves of intestinal DNA viruses at week 1 and 3, respectively. (C and D) Taxonomic composition of intestinal DNA viruses at the family at week 1 (C) and 3(D). (E and F) Taxonomic composition of intestinal DNA viruses at genus level at week 1 (E) and 3 (F), respectively. Data were expressed as the mean (n = 5 or 6 biological replicates).

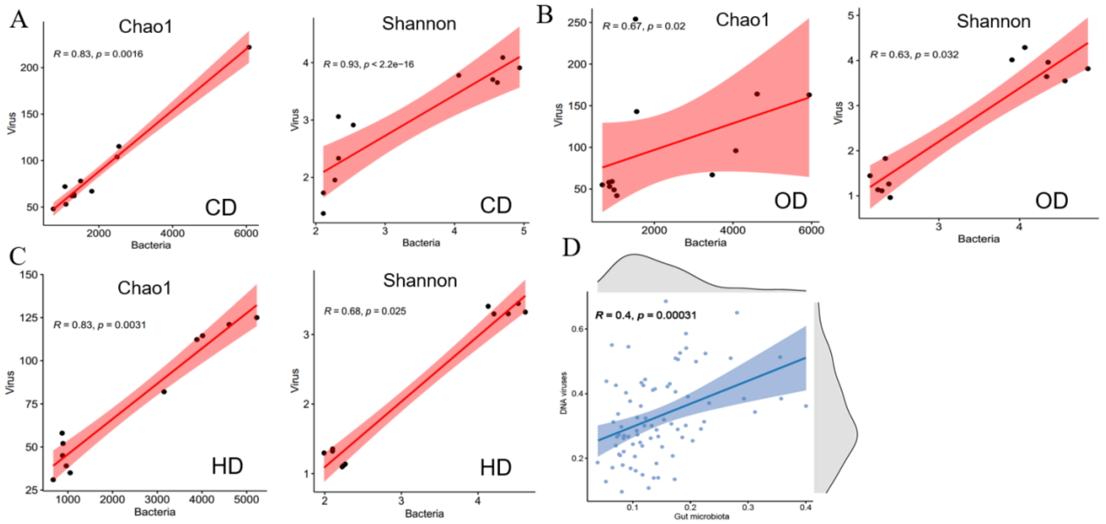


**Figure S.6 Diversity and composition of DNA viruses were associated with the diversity and composition of the gut microbiota**. (A, B and C) Spearman **c**orrelation analysis between intestinal DNA virus and microbiota alpha diversity for samples fed different diets. (D) Spearman correlation analysis between intestinal DNA viral and microbiota alpha diversity for all samples.

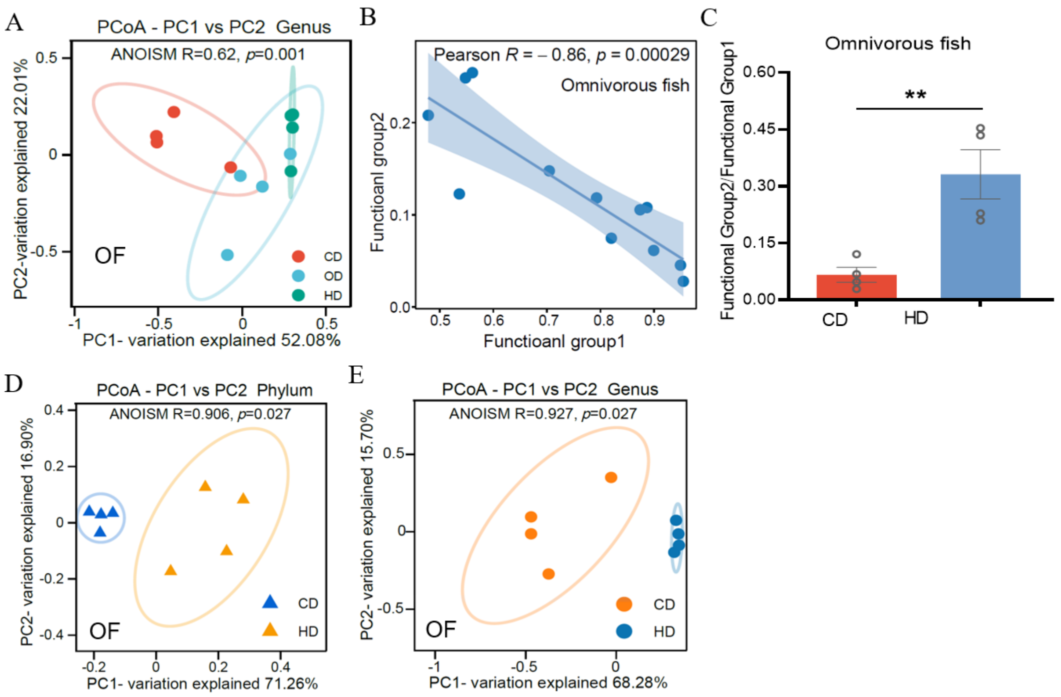


**Figure S.7 The ratio of "Functional Group 2/Functional Group 1" reflects the structural characteristics of the microbiota in omnivorous fish fed CD, OD and HD diets.** (A) PCoA analysis of gut microbiota at the genus level in omnivorous fish (OF, zebrafish) showed that diet altered the gut microbiota. (B) Pearson analysis revealed a significant negative correlation between the relative abundance of functional group 2 and functional group 1. (C) The gut microbiota of zebrafish fed CD and HD diets differed significantly in the ratio of "Functional Group 2/Functional Group 1". (D and E) PCoA analysis the gut microbial communities at the phylum and genus levels. Unpaired t test was used to analyze differences, ** $p<0.01$. Data were from references [15]. Omnivorous fish (zebrafish) were fed with CD and HD diets for two weeks.

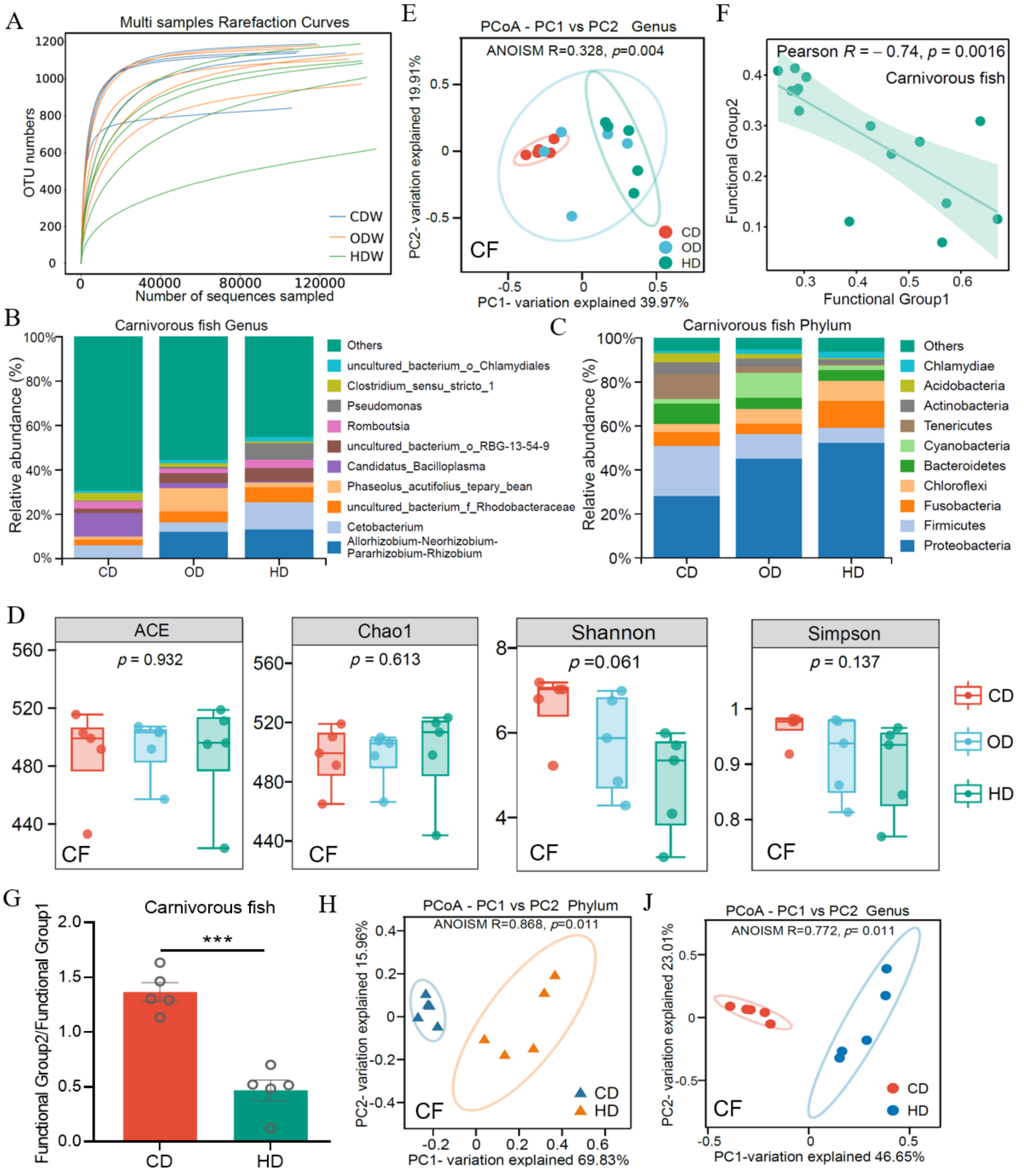

A
Multi samples Rarefaction Curves
OTU numbers
Number of sequences sampled
CDW
ODW
HDW
E
PCoA - PC1 vs PC2 Genus
ANOISM R=0.328, p=0.004
PC2- variation explained 19.91%
PC1- variation explained 39.97%
CF
CD
OD
HD
F
Pearson R = – 0.74, p = 0.0016
Carnivorous fish
Functional Group2
Functional Group1
B
Carnivorous fish Genus
Relative abundance (%)
Others
uncultured_bacterium_o_Chlamydiales
Clostridium_sensu_stricto_1
Pseudomonas
Romboutsia
uncultured_bacterium_o_RBG-13-54-9
Candidatus_Bacilloplasma
Phaseolus_acutifolius_tepary_bean
uncultured_bacterium_f_Rhodobacteraceae
Cetobacterium
Allorhizobium-Neorhizobium-Pararhizobium-Rhizobium
C
Carnivorous fish Phylum
Others
Chlamydiae
Acidobacteria
Actinobacteria
Tenericutes
Cyanobacteria
Bacteroidetes
Chloroflexi
Fusobacteria
Firmicutes
Proteobacteria
D
ACE
p = 0.932
Chao1
p = 0.613
Shannon
p =0.061
Simpson
p = 0.137
G
Carnivorous fish
Functional Group2/Functional Group1
***
H
PCoA - PC1 vs PC2 Phylum
ANOISM R=0.868, p=0.011
PC2- variation explained 15.96%
PC1- variation explained 69.83%
J
PCoA - PC1 vs PC2 Genus
ANOISM R=0.772, p= 0.011
PC2- variation explained 23.01%
PC1-variation explained 46.65%

**Figure S.8 The ratio of "Functional Group 2/Functional Group 1" reflects the structural characteristics of the microbiota in carnivorous fish fed CD, OD and HD diets.** The carnivorous fish (CF, Largemouth bass) were fed a CD, OD and HD diet for three weeks (The diet formulation shown in Table S1)**.** (A) Rarefaction curves of samples in carnivorous fish fed different diets. (B and C) Gut microbial composition in carnivorous fish (Largemouth bass) at the genus and phylum level. (D) Alpha diversity analysis of gut microbiota, including ACE, Chao1, Shannon and Simpson indices. Kruskal-Wallis non-parametric test was conducted to obtain *p*-values for all group differences, followed by Dunn's test to obtain significance levels between the two groups. (E) PCoA analysis of gut microbiota at the genus level. (F) Pearson analysis revealed a significant negative correlation between the relative abundance of functional group 2 and functional group 1. (G) The gut microbiota of Largemouth bass fed CD and HD diets differed significantly in the ratio of "Functional Group 2/Functional Group 1". (H and I) PCoA analysis of the gut microbial community of Largemouth bass fed CD and HD diets at the phylum and genus level. PCoA analysis was performed on all samples using Bray-Curtis's distances and ANOISM was used to analyze variation in microbial communities. Data were expressed as the mean ± SEM (n = 5 biological replicates) and unpaired t test was used to analyze differences in ratios of functional groups, *** $p < 0.001$.

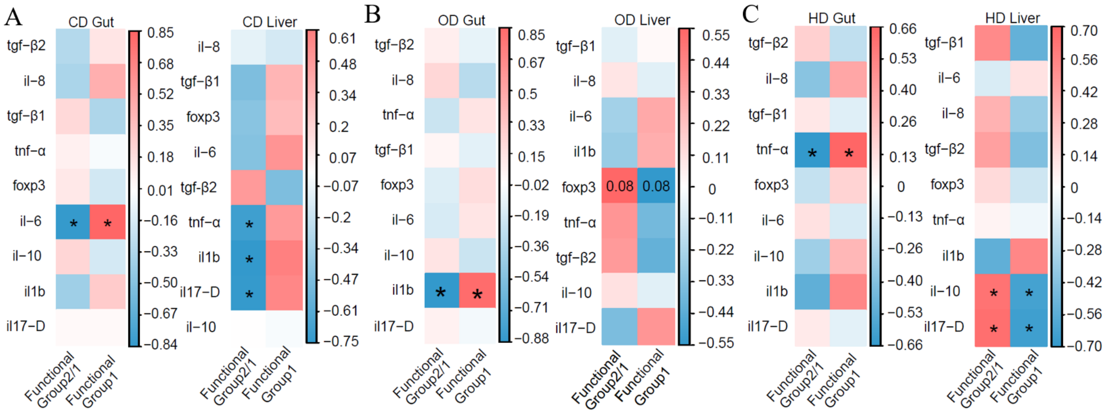


**Figure S.9 The ratio of "Functional Group 2/Functional Group 1" was associated with the host's inflammatory gene expression.** (A, B and C) Spearman correlation between ratio of "Functional group 2/Functional group 1" and mRNA expression levels of host inflammation-associated genes in CD (A), OD (B) and HD (C) dietary groups, respectively. The expression of inflammatory genes was obtained by RNA-seq sequencing. Pro-inflammatory cytokines including il-1β, il-6, il-8, tnf-α and il-17D. Anti-inflammatory cytokines including tgf-β1, tgf-β2 and transcriptional regulatory factor foxp3. il-1β, interleukin-1 beta; il-6, interleukin6; il-8, interleukin 8; tnf-α, tumor necrosis factor-alpha; tgf-β1, transforming growth factor-beta1; tgf-β2, transforming growth factor-beta2; il-10, interleukin10; foxp3, forkhead box P3. The data represented 5 or 6 biological replicates per group, * $p < 0.05$.

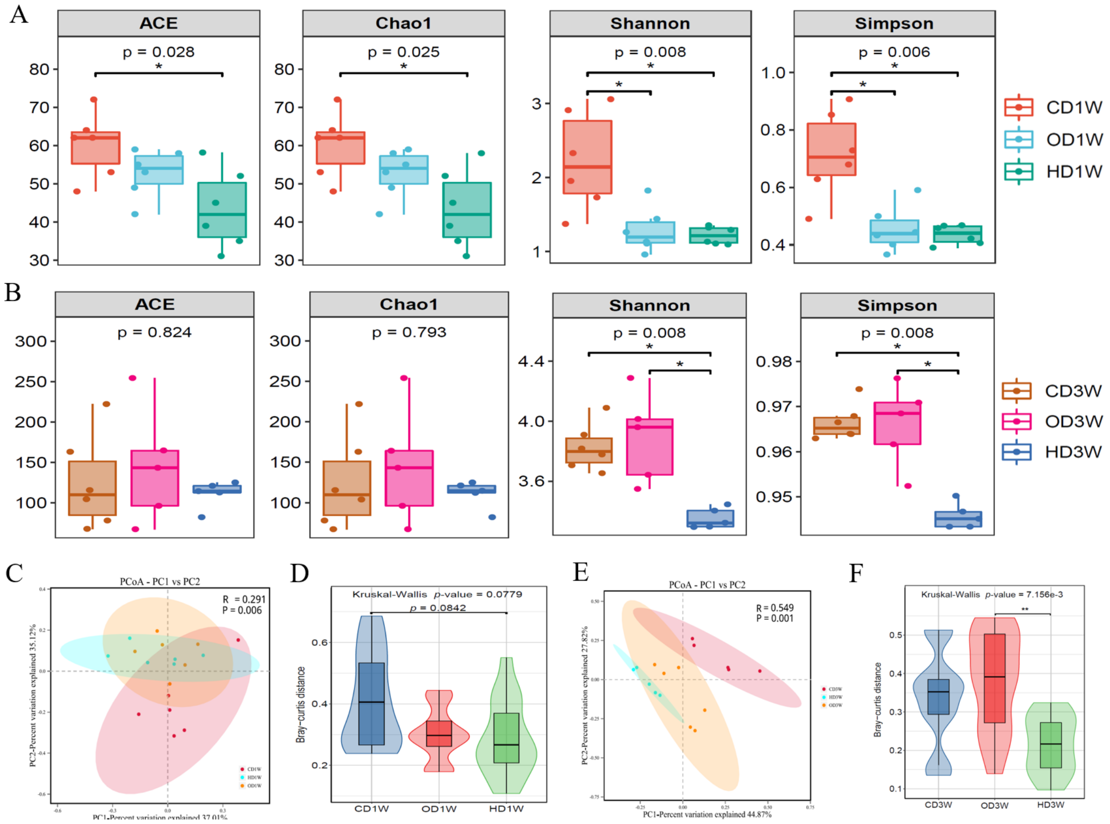


**Figure S.10 The diversity of gut DNA virus.** Alpha diversity of the gut DNA virus at the species level at week 1 (A) and 3 (B), including ACE, Chao1, Shannon and Simpson indices. (C and E**)** PCoA of all samples at the species level by Bray-Curtis's distance. The dotted ellipse borders represent the 95% confidence interval. (D and F) Bray-Curtis's distance was calculated based on grouping distance matrix. Kruskal-Wallis non-parametric test was conducted to obtain *p*-values for all group differences, followed by Dunn's test to obtain significance levels between the two groups. Data were expressed as the mean, * $p< 0.05$ and ** $p<0$ .01(n = 5 or 6 biological replicates).

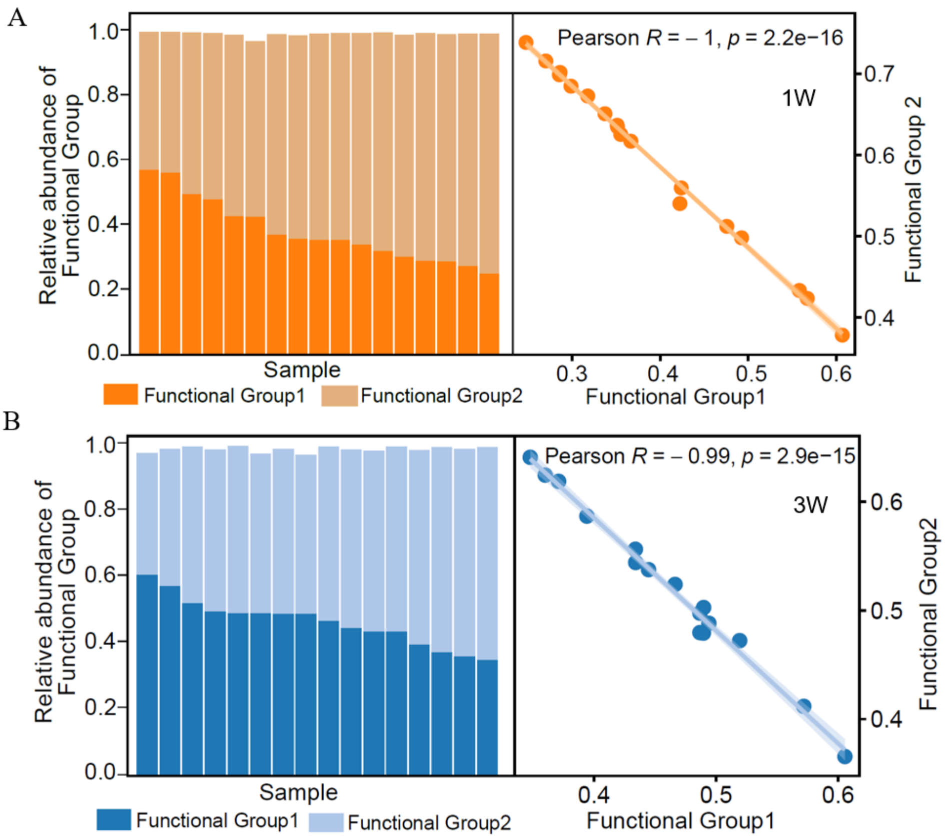


**Figure S.11 Two functional groups show negative correlation in relative abundance.** (A and B) Pearson correlation of the relative abundance of gut microbial Functional Group2 and Functional Group1 at week 1 (A) and week 3 (B) (n = 17 or 16 biological replicates).

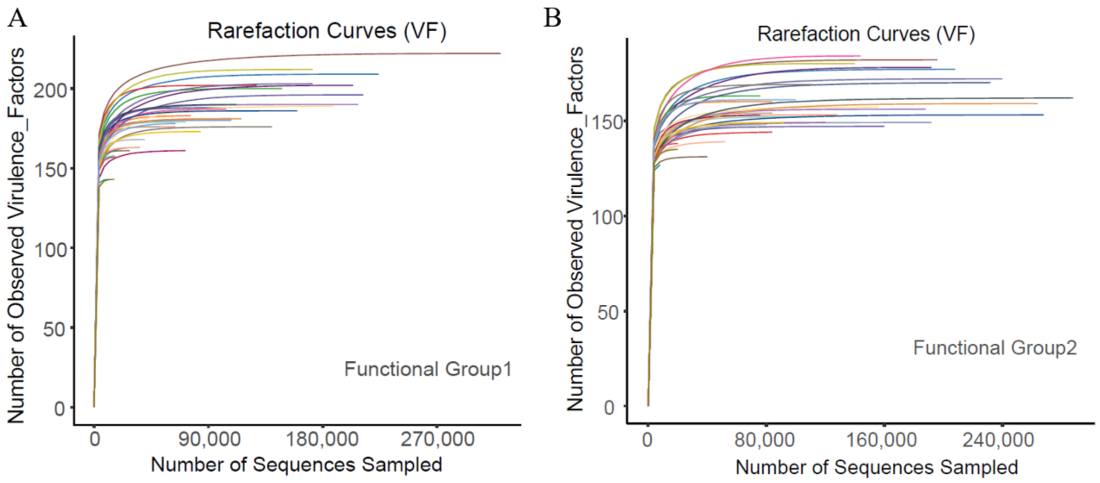


**Figure S.12 Virulence factors gene annotation of two functional groups.** (A and B) Rarefaction curves of virulence factor genes for Functional Group 1 and Functional Group 2 from all samples.
